# *De novo* design of proteins that bind naphthalenediimides, powerful photooxidants with tunable photophysical properties

**DOI:** 10.1101/2024.09.30.615927

**Authors:** Samuel I. Mann, Zhi Lin, Sophia K. Tan, Jiaqi Zhu, Zachary X. W. Widel, Ian Bakanas, Jarrett P. Mansergh, Rui Liu, Mark J. S. Kelly, Yibing Wu, James A. Wells, Michael J. Therien, William F. DeGrado

**Affiliations:** Department of Pharmaceutical Chemistry, University of California at San Francisco, San Francisco, California 94158-9001, United States; The Cardiovascular Research Institute, University of California at San Francisco, San Francisco, California 94158-9001, United States; Department of Cellular and Molecular Pharmacology, University of California, San Francisco, San Francisco, CA 94158, USA; Department of Chemistry, Duke University, Durham, North Carolina 27708, United States

## Abstract

*De novo* protein design provides a framework to test our understanding of protein function and to build proteins with cofactors and functions not found in nature. Here, we report the design of proteins designed to bind powerful photooxidants and the evaluation of the use of these proteins to generate diffusible small molecule reactive species for applications in proximity labeling. Because excited state dynamics are influenced by the dynamics and hydration of a photo-oxidant’s environment, it was important to not only design a binding site, but also to evaluate its dynamic properties. Thus, we used computational design in conjunction with molecular dynamics (MD) simulations to design a protein, designated NBP (<u>N</u>DI <u>B</u>inding <u>P</u>rotein) that held a naphthalenediimide (NDI), a powerful photooxidant, in a programable molecular environment. Solution NMR confirmed the structure of the complex. We evaluated two NDI cofactors in this *de novo* protein, using ultra-fast pump-probe spectroscopy to evaluate light-triggered intra- and intermolecular electron transfer function. Moreover, we demonstrated the utility of this platform to activate multiple molecular probes for protein proximity labeling.

## INTRODUCTION

In nature, proteins bind a diverse array of small molecules to accomplish function from light-harvesting to cellular signaling.^1-2^ These proteins have evolved to recognize molecular shape and electron density distribution, enabling them to bind and tune the properties of organic cofactors within the complex milieu of a cell. The design of proteins that bind small molecules, including functional cofactors is an ongoing challenge^3-4^ often described as that of “inverse drug design”.^5^ Solving this challenge could enable the design of proteins that bind new-to-nature cofactors which leverage the advantages of proteins (genetically encodable, evolvable, expressible) with chemical properties that enable catalysis, photophysics, and photochemistry not found in nature. Synthetic biological photocatalysis (outside of the photosynthetic pathway) has so far been limited to flavoproteins. While flavins perform important functions in nature, they have limited properties when compared to the photocatalysts used in chemical synthesis energy capture and utilization.^6-7^ Alternatively, could we design enzymes for new-to-nature photoredox catalysis using fully *de novo* proteins and inexpensive organic photooxidants? Here, we achieve the goal of designing a naphthalenediimide (NDI)-binding protein with precisely tuned structural and dynamic properties, and demonstrate its utility for photo-initiated proximity-based labeling.

Recent advancements in design methodology spurred by advances in machine learning have enabled the design of proteins that bind organic small molecules and cofactors tightly (low nM to pM).^8-13^ However, until recently, the few successful examples of nM binders have depended on screening of a large library of designs or high-throughput evolution methods^8-9^, ^14-16^ and have been focused on those close to nature or with known binding sites in the PDB.^8-9, 13^ Therefore, the ability to bind non-natural cofactors beyond metalloporphyrins remains challenging. Moreover, for function it is important to not only bind cofactors but also tune their environments including the polarity, water-accessibility, and dynamics of the binding site.

NDIs (Fig. 1) are a well-studied class of easily synthesized molecules with unique electronic, structural, and photophysical properties.^17-18^ Their stability has led to their use in multiple functional materials,^19^ DNA intercalation,^20^ and electron donor-acceptor systems.^21-22^ In addition, NDIs are powerful photooxidants; for example, an NDI bearing no bay region substituents (on the napthyl ring, Fig. 1B) features a singlet excited state reduction potential (^1^E^-/^*) > 2.40 V (vs SCE),^23^ and is thus capable of oxidizing aromatic amino acids Phe, Tyr, and Trp (Fig. 1B).^24^ Combined with the ability to tune their electronic properties through substituent effects and H-bonding interactions, NDIs are enticing targets for protein design^25^ or photosensitizers.^17-18, 26-28^ We study two NDI derivatives that differ in their N-substituents. Phenyl-substituted NDI**1** has a very short-lived singlet state due to intramolecular ET, which creates a charge-separated (CS) state, again with a short ∼ 1 ps lifetime (Fig. 1A)^23^. Alkyl-substituted NDI**2** (Fig. 1B) lacks the pendant phenyl groups and hence does not undergo photoinduced intramolecular ET, instead forming intermolecular CS states with ns lifetimes (Fig. 1A).^29^ The availability of both derivatives allows one to evaluate the importance of lifetime and CS states to influence photophysical and photochemical processes. We initially focused on NDI**1** (Fig 1B) due to its ability to perform intramolecular charge transfer^23^ (Fig. 1A) and its relevance in synthetic donor-acceptor (DA) systems.^17, 30^

**Figure 1.**
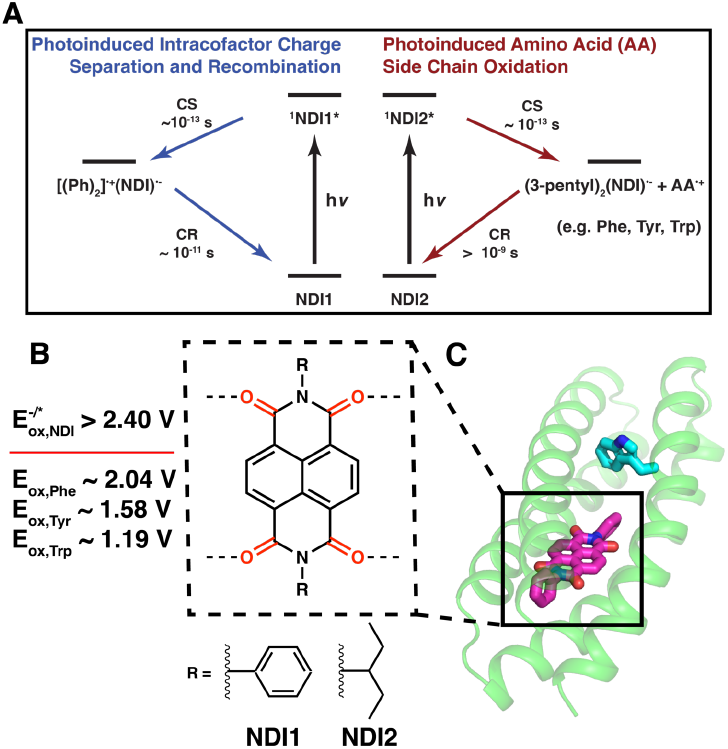
(**A**) Jablonski diagram illustrating photoinduced intramolecular charge separation/charge recombination (CS/CR) (left) and intermolecular CS/CR (right) reactions triggered by photo-excited NDI**1** or NDI2 (^1^NDI*). (**B**) NDI cofactors for intramolecular (NDI**1**) and intermolecular (NDI**2**) photoinduced charge separation reactions. (**C**) General model of an NDI-binding protein used in this study with binding site that includes H-bonds to the carbonyl oxygens (dashed lines). Potentials shown are relative to the SCE reference electrode.

**Figure 2.**
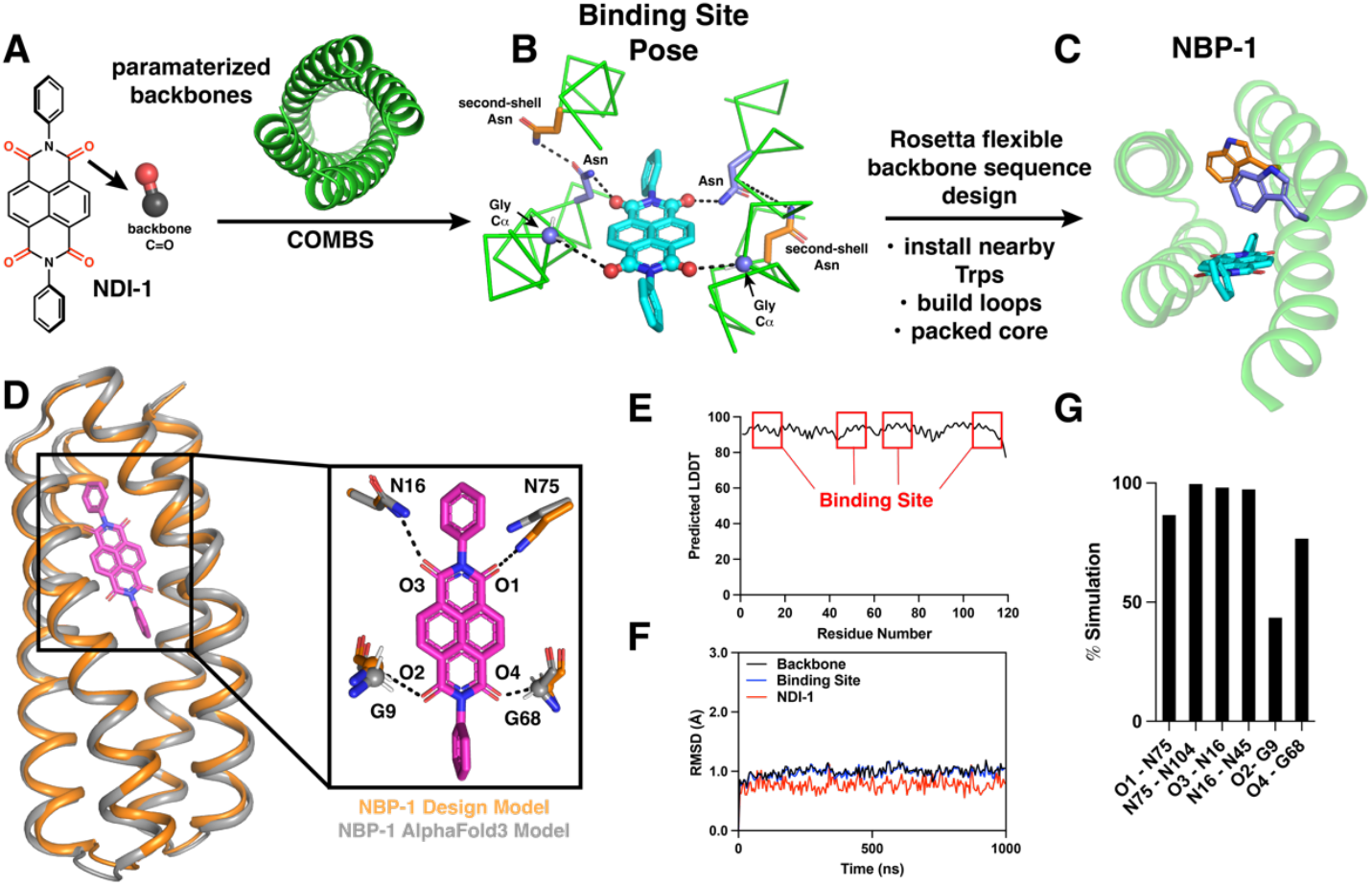
Design process for NBP1. (**A**) NDI carbonyls are approximated with proteinaceous backbone carbonyl FGs in COMBS. (**B**) A library of parameterized backbones are searched using the COMBS algorithm to find a designable pose that provides H-bonding interactions to the NDI (blue residues) and second-shell H-bonding interactions (orange residues). (**C**) The binding site is constrained and Rosetta flexible backbone sequence design is used to design the remaining sequence. Trp residues placed during the design process at 3.5 Å (blue) and 9.5 Å (orange) as possible electron transfer partners for the NDI. MASTER was used to build loops to make NBP1. (**D**) The AlphaFold3 model agreed with the designed model of NBP1 with a helical RMSD of 0.81 Å, including excellent prediction of the designed H-bonding sidechains. (**E**) Plot of pLDDT from AlphaFold3 showing the strong confidence in the designed protein sequence. Red squares indicate residues within 6 Å of the NDI cofactor. (**F-G**) MD simulations helped inform design stability and dynamics. (**F**) Averaged RMSD of NBP1-NDI**1** complex over three 1 µs simulations showing extreme stability of the design and impressive RMSD for NDI**1** cofactor within the binding site. Binding site refers to residues within 10 Å of the cofactor. (**G**) Analysis of designed H-bonding interactions between protein and NDI**1** showing that H-bonds are maintained throughout the simulations

The environment of a protein’s binding site also has a large effect on the electrochemical midpoint potential and excited state lifetimes of cofactors. We sought a system where we could systematically vary the polarity, water-accessibility, and dynamic behavior of the binding site to determine their effects on complex photo-oxidative functions.^31^ The design of proteins that simply bind small molecules without additional considerations of building a functional environment and dynamics has not been accomplished for cofactors other than metalloporphyrins, where metal ligation thermodynamically stabilizes the complex while also providing rigid geometric restraints. Indeed, design of proteins that merely bind small molecules rich in polar substituents has been described as a grand challenge in protein design.^32^

Here, we sought to bind NDI derivatives in a fully solvent-inaccessible environment, while providing sites to engage their four polar carbonyl groups. We accomplish this task using a computational method that focuses attention on the key binding interactions (including second shell H-bonds) through a computational construct referred to as a van der Mer (vdM). VdMs define favorable positions to bind functional groups such as carbonyls and aromatic rings relative to the backbone of a designed model. We found this approach to be particularly useful when combined with other computational methods, once the key residues were placed on a candidate backbone. We then use MD to evaluate the structural stability of the NDI in the binding site and its sequestration from solvent.

Using these methods, we designed a set of three NDI binders, designated NBP1-3 (NDI Binding Protein), all of which bound NDI**1**. The highest affinity design (NBP1) bound NDI**1** with 27 nM affinity, and its structure was validated by solution NMR. We then evaluated the photophysical properties using fs to ns laser spectroscopy, which illustrated the influence of the pendent N-substituents and the cofactor’s environment on CS lifetimes. We also demonstrate their ability to perform photo-triggered intermolecular electron transfer to activate reactive probes in solution for proximity protein labeling.

## RESULTS

### Design of NBP1 and MD simulations

Initially we targeted binding of NDI**1** (Fig 1B) due to its ability to form intramolecular charge separate states and its N-phenyl substituents which are advantageous for good packing with amino acid sidechains. While the NDI derivatives discussed here are *D*_*2*_ symmetric, they can be derivatized to be asymmetric.^33^ Taking advantage of this symmetry offers an excellent starting point by parameterizing *D*_*2*_-symmetric four-helix bundles that can then be expanded to any NDI scaffold and de-symmetrized during flexible backbone sequence design. In addition to the asymmetric environment provided by the side chains, the twist of the coiled coil can differentiate prochiral centers, providing an approach to modulate the spectroscopic and electronic properties of the NDI in a way that would be difficult to do outside of a protein. Using COMBS, we generated ligand poses on a small library of parametrized helical bundles^34^ that included classic as well as C*α*-H H-bonds, illustrating the versatility of this method. The carbonyl groups on the NDI were approximated with amino acid backbone carbonyls^9^ (Fig. 2A, see SI), which were used to search for polar interactions to the ligand. In the most promising pose two Asn side chains and two Gly C*α* each donate a single H-bond to one of the four NDI carbonyl groups each donating an H-bond to a carbonyl oxygen, as well as two Gly-C*α* donating H-bonds to the remaining two carbonyl oxygens (Fig. 2B). Gly Cα-H H-bonds, while weaker than prototypical polar H-bonds, are important in ligand binding^35^ and enforcing secondary structural motifs,^36^ and are thus motifs that should be adequately sampled. In addition, we implemented a recursive version of the COMBS algorithm^8^ which allowed us to find vdMs that provide second-shell H-bonding interactions to re-enforce the binding site. With this strategy, we found second-shell Asn residues that donated H-bonds to the carbonyl oxygens of the first-shell Asn donors (Fig 2B). This binding site was constrained during the remaining sequence design (see SI).

Because NDIs possess substantial ^1^E^-/^* values, Trp residues were placed at 3.5 Å and 9.5 Å from the NDI (measured closest approach to naphthalene core) as possible electron donors for photoexcited NDI. In addition, there were three Phe residues within 10 Å of the NDI. The full sequence design was accomplished using a custom RosettaScript for flexible backbone sequence design in conjunction with the MASTER algorithm^11, 37^ (See SI). The resulting sequence, designated NBP1 has no similarity with any natural proteins (blast *E* value One challenge in ligand binding is accommodating polar residues, such as the four carbonyl groups of NDI, within the apolar core of a protein. This often leads to instability and increased dynamics of the protein backbone and sidechains. Therefore, we wanted to assess the use of molecular dynamics (MD) as a design tool to help predict successful designs and stable binding site-cofactor interactions. We next conducted three independent 1 µs MD simulations of the resulting model to probe its conformational stability and dynamics in the presence of water. These properties have been shown to be important in tuning photophysical function.^38^ The simulated ensembles can then be readily assessed and validated with solution NMR spectroscopy (*vide infra*). Overall, the designs showed impressive stability throughout the simulations with average backbone RMSDs all below 1.15Å (Figs. 2F, S2). While the protein backbone and binding site in NBP1 were very stable (average RMSD of 0.99 Å and 0.97 Å, <0.05 against the nonredundant protein database). Two additional designs (NBP2-3) were prepared from the same backbone and had the same NDI H-bonding interactions but contained varying aromatic residues near the NDI binding site (Fig. S1). respectively), the NDI**1** was exceptionally stable (average RMSD 0.77 Å), illustrating the precisely designed binding site. Moreover, the designed H-bonding interactions were present with >85% occupancy over the simulation (Fig. 2G), indicating the four carbonyls were well accommodated within the protein core. The NDI remained highly dehydrated throughout the simulation (Fig. S3); indeed, the entire protein interior remained dry with the exception of three small pockets of water located at least 4.5 Å from the closest heavy atom of the ligand. Note that these waters a not within H-bonding distance to NDI-carbonyls.

For additional confirmation of the structural stability of apo-NBP1, we ran *ab initio* folding predictions in Rosetta^39^ and compared the outputs to AlphaFold3^40^ structure predictions. The *ab initio* calculations showed excellent agreement with our design where the top scoring structure had a backbone RMSD = 0.7 Å (Fig. S4) The AlphaFold3 predictions also confidently predicted the overall structure with an average pLDDT (predicted local distance difference test, a per-residue measure of local confidence) 88 and all backbone RMSD of 0.81 Å (Fig. 2D-E).

### NDI1 binds cofactor with low nM affinity in a rigid, apolar binding site

All designed proteins express well in *E. coli* (∼20-30 mg/L) and were purified to homogeneity. Spectral titrations indicate that NBP1-3 bound NDI**1** with similar shifts in the absorbance spectrum (Figs. 3A, S5). While all three designs bound the cofactor, NBP**1** had the best solubility properties and was chosen for further characterization. Despite the cofactor’s extremely poor water solubility and inherent disposition towards aggregation, NBP**1** tightly binds one equivalent of NDI**1** (Fig. 3B). NDI**1** was added from a DMSO stock solution (binding occurred within the mixing time) and resulted in a shift of the absorption bands that was characteristic of an apolar binding site. A spectral titration of the change in absorption as a function of the protein concentration is well described by a single-site binding isotherm with an apparent dissociation constant of 27 nM (*vida infra*).

**Figure 3.**
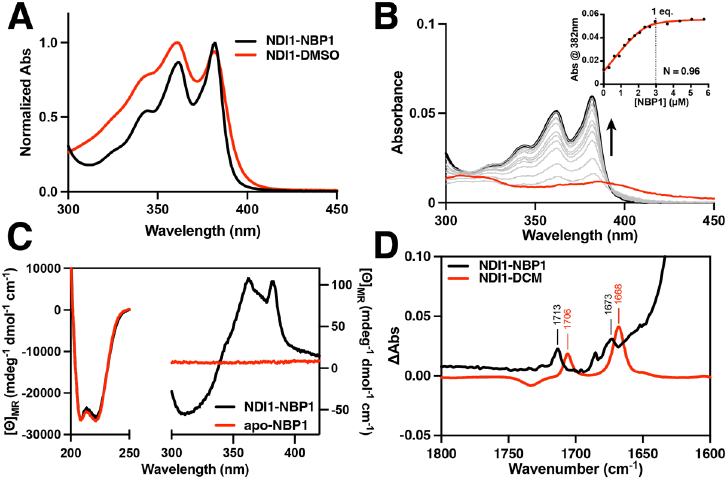
(A) Electronic absorption spectra of holo-NBP1 and free NDI**1** in DMSO showing a shift in the absorbances indicative of binding. (**B**) UV-vis titration of apo-NBP1 into a solution of NDI**1** (3 µM) giving a 1:1 stoichiometry. (**C**) CD spectra showing the helical structure of apo- and NDI**1**-NBP1 and the induced Cotton effect for the NDI absorbances at 362 and 382 nm in the visible region. (**D**) IR difference spectra showing a shift in the C=O vibrations between free NDI1 in methylene chloride solvent and NDI1 in NBP1 due to designed H-bonding interactions.

NDI**1** exhibits a weak, broad absorbance at 388 nm (Fig. 3A-B, red) when added to aqueous buffer due to aggregation.^41^ Once isolated within the protein’s binding site the bands split, revealing characteristic NDI vibrational fine structure (343, 362, and 382 nm) with a measured extinction coefficient (ε_382_= 19,100 M^-1^cm^-1^) consistent with published values in organic solvent, indicating the NDI**1** is held rigidly in the binding site.^23, 33^ Similarly, while NDI**1** is achiral, we expect placement of this synthetic cofactor within the designed rigid binding site to induce chirality if it was rigidly held in a chiral environment. Indeed, the visible CD spectrum of holo-NBP1 shows a clear induced Cotton effect associated with the absorption bands at 362 and 382 nm (Fig. 3C).

To determine the effect of the designed H-bonding interactions on the C=O vibration of the NDI, we measured solution IR spectra of the NDI**1** and holo-NBP1. We expected to observe the NDI C=O vibrations around 1650 cm^-1^,^42^ however they were obscured by the intense amide I stretching frequency from the helical backbone (∼1600-1700 cm^-1^).^43^ To address this, we expressed ^13^C-^15^N labeled NBP1 to shift the amide I frequency to ∼1590 cm^-1^,^44^ allowing us to better view the NDI C=O vibrations. In so doing, we saw a shift in the C=O vibrations between free NDI**1** in DCM and holo-NBP1 (Fig. 3D). The dielectric of DCM has been shown to be similar to the apolar environment of a protein interior, thus is a reasonable estimate for a binding site void of H-bonding interactinos.^30^ In particular, the band seen at 1706 cm^-1^ for the free NDI was clearly separated from the amide absorption of the protein, and this vibration moved to 1713 cm^-1^ once bound to the protein, indicative of a change in the environment surrounding the cofactor.

### Contributions of H-bonding interactions to the affinity of NBP1 for NDI

To probe the contribution of H-bonding to the free energy of binding of the cofactor, we determined the apparent dissociation constant for binding of NDI**1** to NBP1 and a mutant in which the first-shell Asn residues were converted to Ala (N16A/N75A, Fig. 4A). NBP1 has two Trp residues within 10 Å of the cofactor, which is well within the expected Forster distance for energy transfer. Indeed, we observed a quenching of the Trp fluorescence in holo-NBP1. A fluorescence titration of NBP1 with NDI**1** showed a linear increase in the degree of quenching until a stoichiometry of 1:1 is reached, after which no further quenching was observed. This behavior is consistent with relatively tight binding, making it challenging to obtain an accurate dissociation constant from a single titration experiment. To improve accuracy, we conducted a series of four titrations at four different total protein concentrations, demonstrating a classical binding isotherm; global non-linear least squares fit of the full set of data provided an apparent dissociation constant of 27 ± 6.7 nM (Fig. 4C). As expected, removal of the two H-bonding Asn residues led to a marked decrease in binding affinity. A titration of NBP1-N16A/N75A indicated that this mutant had a dissociation constant of 470 nM corresponding to a 1.7 kcal/mole decrease. The change in the H-bonded environment between NBP1 and the double-Asn mutant (N16A/N75A) also leads to a change in the cofactor’s electronic absorption spectrum indicating that the electronic structure of NDI is sensitive to the environment (Fig. 4B), consistent with previous studies of the effect of solvent polarity on the spectrum of NDI derivatives.^45^

**Figure 4.**
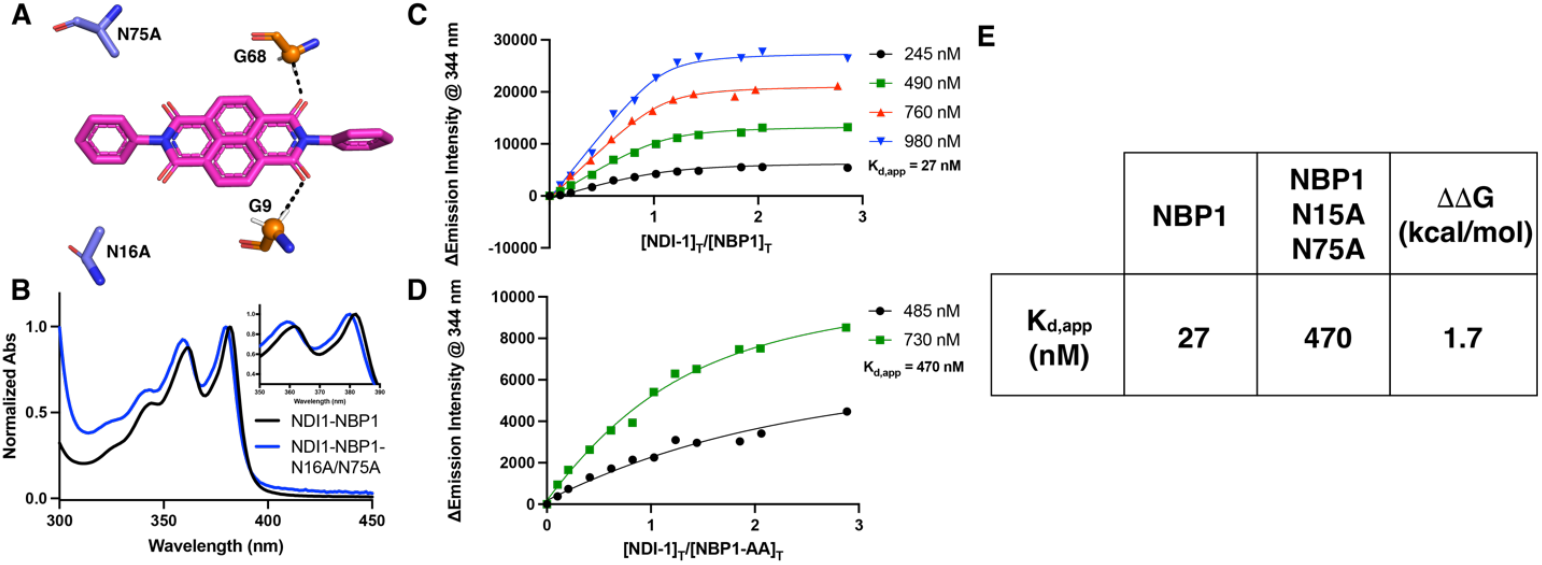
(**A**) Model showing Asn to Ala mutations made in NBP1 to disrupt H-bonding interactions. (**B**) Electronic absorption spectra showing shift in absorbance due to modification of H-bonding environment. Inset: Zoom of absorption maxima. (**C**-**D**) Tryptophan fluorescence quenching experiments showing the concentration dependent change in Trp emission at 344 nm. (**E**) Apparent Kd values from fits of C and D, and calculated ΔΔG value associated with the loss of two H-bonds to NDI**1**.

### Structural validation using NMR spectroscopy

Solution NMR confirmed our designs and corroborated the MD simulations. Free NDI1 in d6-DMSO showed a single 1H signal for the four naphthalene protons, consistent with the D2 symmetry of this cofactor (Fig. S6A). However, within the protein, NDI1 is expected to be bound in an asymmetric environment in which each naphthalene proton is in a unique magnetic environment. Indeed, the chemical shifts show that the NDI protons become non-equivalent in the complex, providing more evidence for the precise placement of the cofactor (Fig. S6B). The sidechains of the protein also were very well dispersed in the complex, and many of the resonances of the binding site residues signals are shifted extremely upfield by as much as 2 ppm due to the strong ring-current above and below the NDI naphthalene core (Table S1, Fig. S7). These data show that the protein has a well-defined structure that positions the NDI in a single orientation within the binding site.

We used three-dimensional NMR to evaluate the structure of the complex. 13C-15N labeled NDI1-NBP1 displayed an exceptionally well-resolved 1H-15N HSQC NMR spectra (Figs. S7-S8), congruent with a unique, well-defined structure. The spectral assignments were obtained for 92.3% of the backbone and 69.3% of the sidechain resonances (see SI). These chemical shifts were input into CSRosetta^46-47^ to compute a model of its tertiary structure (Fig. 5A). The resulting model (PDB 9DM3) matched the design exceptionally well (helical backbone RMSD 0.9 Å ± 0.18 Å) confirming the unique structure observed by MD. Moreover, we measured 19 nuclear Overhauser effects (NOEs) between NDI1 and the neighboring sidechains (Fig. 5B, Table S2). Each of these NOEs are in very good agreement with the corresponding proton-proton distances observed in the weighted ensemble of structures computed in our MD simulations (Figs. 5C, S9), which showed that the corresponding protons were within 5 Å. In summary, the unrestrained MD simulations, which were conducted using the design model as a starting structure, were in excellent agreement with the overall backbone structure computed by CSRosetta as well as the detailed contacts reflected in the protein-NDI NOEs (Fig. 5C, Table S2). This illustrates the effectiveness of using MD as a guiding tool during the design process.

**Figure 5.**
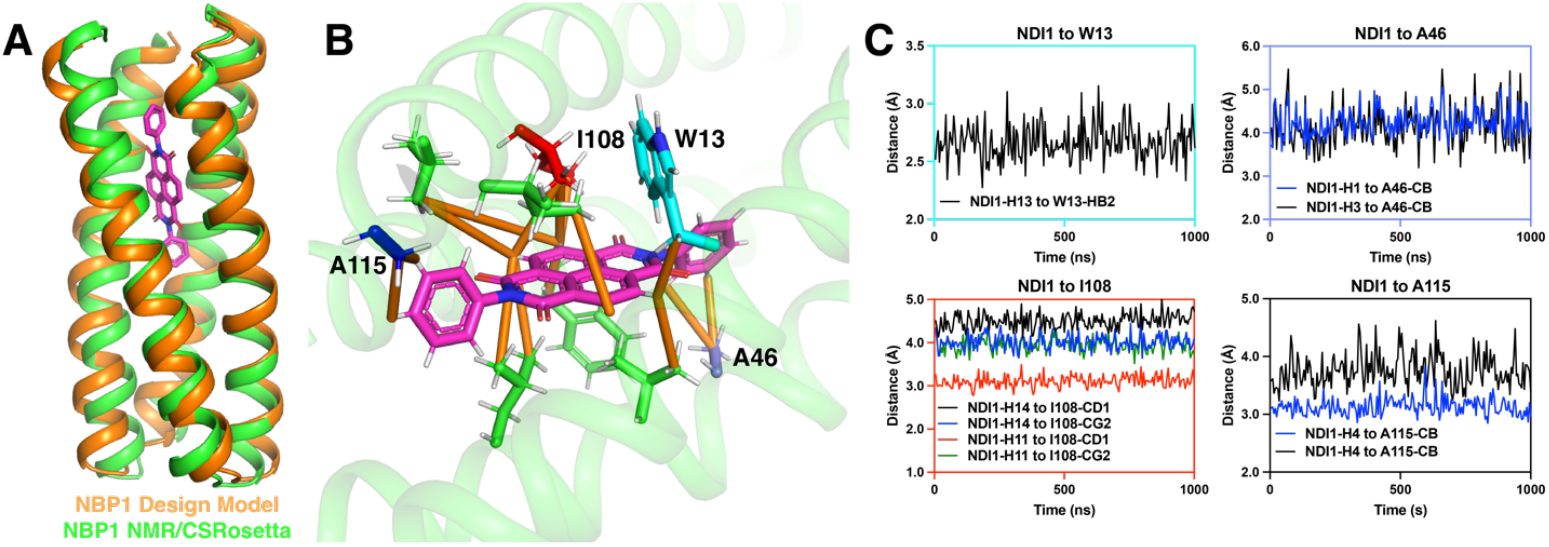
(**A**) Overlay of NDI**1**-NBP1 design model (orange) and calculated NMR structure using CSRosetta (green, PDB 9DM3) showing 0.9 Å helical backbone rmsd. (**B**) 17 protein-NDI NOEs (orange lines) experimentally determine the location and orientation of the NDI**1** cofactor within the binding site of NBP1. (**C**) Analysis of the MD simulations of NDI**1**-NBP1 allow us to corroborate measured NOEs. Plots show atom-atom distances over the course of the MD trajectory (see SI for additional detail). All measured NOEs maintained atom-atom distances of <5 Å. Plot boxes are color coded to match their respective residues in (**B**).

### NDI Photophysical Properties

Photophysical characterization of NDI**1**-NBP1 and NDI**2**-NBP1 was carried out using pump-probe time-resolved transient optical methods. Electronic excitation (λ_ex_= 380 nm) of the cofactors in these proteins produces the potent ^1^NDI* photo-oxidant. Congruent with data acquired in solution, because the NDI**1** cofactor features imide phenyl (Ph) substituents (Figs. 1B, S10), and a ^1^NDI* state ^1^E^-/*^ potential having sufficient thermodynamic driving force to oxidize unsubstituted arenes,^23^ an ultrafast photoinduced intramolecular CS reaction is observed for NDI**1**-NBP1 [^1^(Ph)_2_NDI* → [(Ph)_2_]^•+^ (NDI)^•-^]. Femtosecond transient absorption (fs-TA) spectroscopic data (**Fig. 6A**) highlight the ^1^NDI* S_1_→ S_n_transient absorption signal; note that its decay is correlated with the rise of another transient absorption signal at 480 nm, corresponding to the radical anion NDI^•-^.^48-50^ These dynamical data characterize the time constant for photoinduced intramolecular CS in NDI**1**-NBP1 (τ_CS_< 1 ps). Decay of the NDI^•-^ transient signal tracks the dynamics of the intramolecular thermal charge recombination (CR) reaction ([(Ph)_2_]^•+^ (NDI)^•-^] → (Ph)_2_NDI; τ_CR_∼ 50 ps).

**Figure 6.**
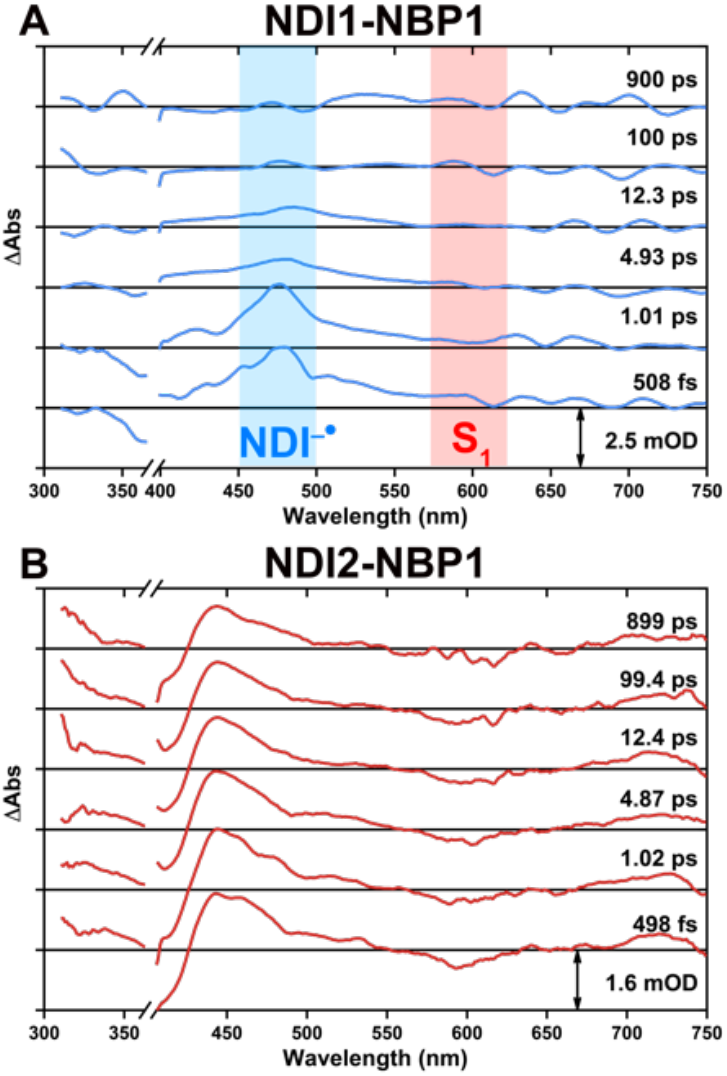
Pump-probe transient dynamics of NBP1 bound to (**A**) NDI**1** displaying intramolecular charge separated species NDI^-•^ (blue) forming as the S1 excited state absorption (red) decays and NDI**2**. Experimental conditions: Buffer solvent system; ambient temperature; NDI**1**, λ_ex_= 385 nm; NDI**2**, λ_ex_= 385 nm.

Because NDI**2**-NBP1 features a cofactor in which the NDI N-phenyl substituents are replaced with N-(3-pentyl) groups, a light-triggered intramolecular charge transfer reaction is not possible. Transient absorption spectroscopic data highlight that following optical excitation of NDI**2**-NBP1, the NDI^•-^ transient absorption persists over the 500 fs - 1 ns time domain chronicled in **Fig. 6B**. These data are consistent with a photoinduced intermolecular CS reaction that likely involves oxidation of a nearby amino acid (AA) side chain (^1^(3-pentyl)_2_NDI* + AA → (3-pentyl)_2_NDI)^•-^ + AA^•+^) such as Trp, followed by subsequent hole (positive charge) migration that transfers this oxidizing equivalent to a more distant AA site; in the NDI**2**-NBP1 protein, the timescale of CR exceeds that which may probed in this experiment (> 7 ns).

### Photo-triggered protein labeling

These NDI**2**-NBP1 photophysical properties indicate its potential utility for phototriggered protein labeling (**Fig. 7A**).^28, 51-53^ Proximity labeling proteomics (PLP) methods are able to spatially label and identify nearby proteins in complex cellular environments by generating reactive intermediates of varying lifetimes .^28^ These tools have provided important understanding of protein interactomes within cells.^52^ Our NDI binders are ideal for this application, with NDI**2** being particularly promising as electronic excitation drives migration of an oxidizing equivalent to distant protein sites. Currently, four types of labeling probes have been used widely in the proximity labeling field^28^ including diazirine (P1), aryl-azide (P2), hydrazide (P3), and phenol-biotin (P4) (Figure S11A). ^54-57^ Diazirine, aryl-azide, and phenol based probes function by formation of reactive carbene, nitrene, and phenol radical species, respectively, via activation by a photocatalyst after irradiation with bio-compatible blue or green light. Alternatively, hydrazide-based probes rely on activation of singlet oxygen (by a photooxidant) to oxidize surface residues on a protein before subsequent covalent modification by the hydrazide moiety. These probes form intermediates with lifetimes ranging from ns to >100 µs, thus determining how far they can diffuse and spatially differentiate nearby proteins.

**Figure 7.**
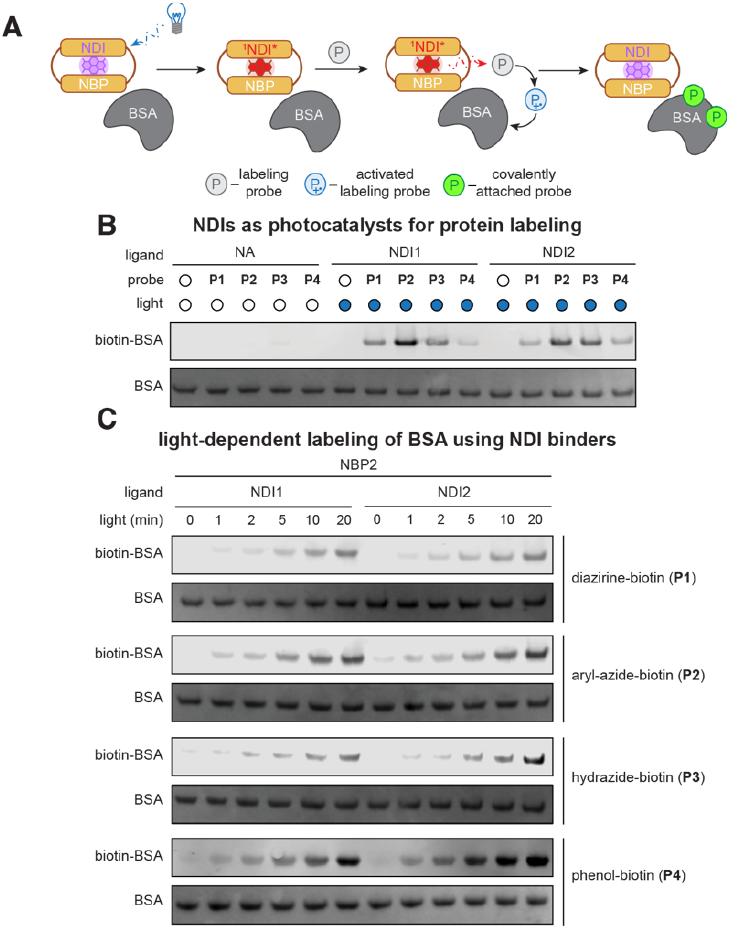
NDI binders perform photo-triggered protein labeling. (**A**) Schematic illustrating NDI binders labeling BSA in solution in a light-dependent manner. (**B**) Western blot of light induced labeling of BSA with free NDI cofactors and (**C**) NBP2 binder with both NDI cofactors showing their ability to activate and label BSA with four labeling probes.

Current methods, however, rely on covalent attachment of an activated photocatalyst (such as dibenzocyclooctyne (DBCO)-Eosin Y (EY)) via azide-functionalized antibodies.^28^ This covalently modified antibody is then used to bind to a protein on the surface of a cell to reveal what is in the neighborhood of that protein. This method is limited by heterogeneous labeling of the antibody and access to only extracellular proteins. Could we instead use NBP1 as a genetically encodable protein that could ultimately be tagged to any protein of interest, rather than relying on the availability of effective antibodies?

To test whether our designed NDI-binders were competent for this type of labeling strategy, we first assessed which NDI derivative was most competent for activation of the labeling probes. We confirmed that, upon illumination with LED light, both free-NDI compounds (NDI**1** and NDI**2**) can activate all four probes to label bovine serum albumin (BSA) (**Fig. 7B**). We found that P2 (the aryl-azide probe) demonstrated the highest level of labeling on BSA with free NDI cofactors, therefore we used P2 to screen NDI binders to ensure reactivity is retained when bound to the protein (Fig. S11C). After confirming that no significant change in the absorption spectrum was observed after irradiation of all the binders (Fig. S11B), we screened three NDI binders (NBP1, NBP2, and NBP3) with both NDI cofactors for self-labeling (Fig. S11C). We found that NBP2 demonstrated the best self-labeling efficiency (compared with NBP1 and NBP3), with slightly enhanced performance for NDI**2** relative to NDI**1**. This was not surprising given the NDI**1** proclivity for intramolecular charge transfer, which is avoided in the alkyl-NDI derivative. Since NBP2 exhibited the best performance, we then tested NDI**1**-NBP2 and NDI**2**-NBP2 to perform photolabeling on BSA using P1-P4 to gauge the ability to activate a range of probes. By performing time-dependent labeling reactions, we observed a significant accumulation of biotinylated BSA with all probes (Fig. 7C). BSA labeling was also light-dependent, reaching a plateau with 10-20 min of irradiation. These results demonstrate that NDI binders are capable of intermolecular electron transfer to labeling probes in solution and can label proteins successfully *in vitro*. Moreover, the modulation in activity between protein scaffold and NDI-derivative suggests this could be a good platform for further development of genetically encodable protein labeling tools.

## DISCUSSION

The ability to bind diverse ligands with high affinity opens up a world of function beyond binding, including catalysis, photophysics, and photochemistry. In this study we augmented our protein design methods with MD to build polar binding sites for abiological NDI cofactors. Using COMBS, we built a binding site that included four H-bonding interactions to the NDI (traditional H-bonds, as well as Gly Cα H-bonds) and designed three proteins that all bound the desired cofactor, including one that bound with ∼30 nM affinity. This speaks to the power of our design process where we can design successful binders without the need for high-throughput experimental screening or evolution. In addition, the use of solution NMR allowed us to readily assess the success of our designs. We identified a substantial number of NDI**1**-NBP1 NOEs which were corroborated with our MD simulations and showed that the calculated backbone structure of holo-NBP1 matched the design with 0.9 Å RMSD. While we used 1µs MD simulations to assess solution dynamics and hydration of the cofactor, an analysis of shorter runs indicated that 200 ns simulations are likely sufficient for rapid assessment directly from design models. This also allows for explicit consideration of water molecules, which are important in ligand binding as well as controlling the polarity and, therefore, photophysical properties of bound molecules. In addition, we utilized a new structure prediction tool, Chai-1^58^, which allows for structure prediction of ligand-protein complexes (see SI). Due to the recent release of this tool, it was not trained on the NBP1 sequence. Chai-1 predicted a NDI**1**-NBP1 structure that matched the designed backbone with 1.07 Å RMSD and predicted all buried side-chains with 1.45 Å RMSD (Fig. S14).

NDIs are powerful photooxidants with tunable photophysical properties. Using fs-TA, we showed that NDI**1** bound to NBP1 was able to form the potent ^1^NDI* photo-oxidant and undergo photoinduced intramolecular CS, forming a short lived (<1 ps) charge-separated state [^1^(Ph)_2_NDI* → [(Ph)_2_]^•+^(NDI)^•-^]. NBP1 also included oxidizable Trp residues in close proximity (3.5 and 9.5 Å), poised to undergo intermolecular electron transfer with a photoactivated NDI. Therefore, we employed the synthetic tunability of the NDI scaffold and replaced the N-phenyl groups with N-3-pentyl substituents (NDI**2**) to avoid intramolecular photo-triggered oxidation. As a result, we observed a photoinduced long-lived (>7 ns) CS resulting from intermolecular electron transfer between NDI and, likely, an oxidizable AA residue (^1^(3-pentyl)_2_NDI* + AA → (3-pentyl)_2_NDI)^•-^ + AA^•+^).

Finally, to illustrate the potential for this system we showed the ability of the designed NDI binders to activate four labeling probes in a light-dependent manner and label BSA *in vitro*. This platform shows great potential to harness and tune the photophysics of bound NDIs by leveraging protein design with cofactor synthetic modification for use in photochemical applications.

## ASSOCIATED CONTENT

### Supporting Information

The Supporting Information is available free of charge on the ACS Publications website.

Methods for protein design, protein expression, NMR, MD, spectroscopy, and photolabeling experiments, along with Figures S1-S13, Tables S1-S2, and Scheme S1 (PDF)

### Accession Code

NMR Chemical Shift information has been deposited in Biological Magnetic Resonance Data Bank (BMRB) with accession number 52600. CSRosetta structure has been deposited in the Protein Data Bank with accession code 9DM3.

## AUTHOR INFORMATION

## ACKNOWLEDGMENT

The authors acknowledge the National Science Foundation (CHE-2109020; NSF CHE-2108660), the Keck Foundation, and National Institute of Health (GM R35 122603, 1R01CA248323-01; 5K99GM143529-02; 1S10OD023455 (UCSF NMR core facility) and PBBR TMC award (UCSF NMR core facility)) for supporting this work. MJT is indebted to the United States Department of Energy (DE-SC0001517) and the United States Air Force Office of Scientific Research (FA9550-20-1-0121) for research infrastructure enabling transient dynamical spectroscopic experiments.

## Insert Table of Contents artwork here

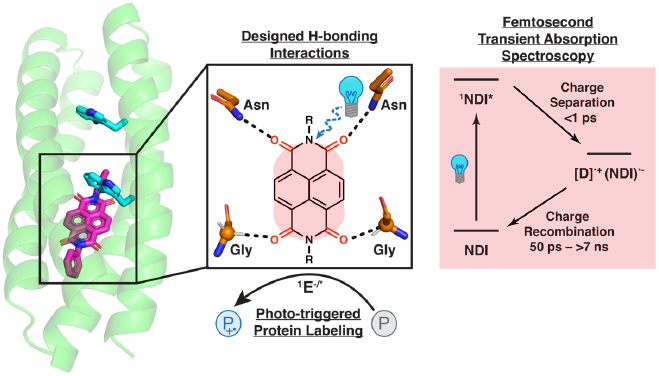

