## Supplemental Information for "*De novo* design of proteins that bind naphthalenediimides, powerful photooxidants with tunable photophysical properties"

**Contents**

| | |
|---|---|
| **Methods** | **S1-S7** |
| **Scheme S1** | **S7** |
| **Fig. S1** | **S8** |
| **Fig. S2** | **S9** |
| **Fig. S3** | **S9** |
| **Fig. S4** | **S10** |
| **Fig. S5** | **S10** |
| **Fig. S6** | **S11** |
| **Table S1** | **S11** |
| **Fig. S7** | **S12** |
| **Table S2** | **S13** |
| **Fig. S8** | **S14** |
| **Fig. S9** | **S15** |
| **Fig. S10** | **S15** |
| **Fig. S11** | **S16** |
| **Fig. S12** | **S17** |
| **Fig. S13** | **S17** |
| **Fig. S14** | **S18** |
| **Fig. S15** | **S18** |
| **Rosetta Code** | **S19** |
| **References** | **S29** |

### Methods

**NBP-1 Design Process.** The design of NBP-1 began with a small library of parameterized 4-helix bundles. [1] The $Ph_2$NDI (NDI-1, Fig. 1B) cofactor binding site was found by running COMBS[2-3] on the library of backbones and analyzing the outputs for maximum number of designable H-bonds (the carbonyl groups of the NDI were approximated by proteinaceous backbone carbonyl groups). Only buried residues were considered, and these were filtered using an alpha-hull algorithm in COMBS, which defines the surface of the protein from its backbone coordinates. The most designable binding site included H-bonds to all four carbonyl groups of the NDI, including two Asn H-bonds and two Gly Ca H-bonds. Moreover, we employed a recursive version of the COMBS algorithm[2] and found second-shell Asn that donate H-bonds to the first-shell Asn H-bond donors. A flexible backbone sequence design protocol[4] was used (see below) to design the backbone sequence and pack around the NDI cofactor, while constraining the first- and second-shell H-bonding interactions.

As previously reported, loops were selected to generate a single chain protein using MASTER.[4-5] Briefly, segments of adjacent helices (6 residues from each helix) were queried against a database of structures from the PDB. Using the "wgap" option in MASTER, loops of specified length connecting these helices were found within a given rmsd cutoff (generally <1.5 Å). These loops were then clustered on rmsd and scored based on number of outputs in a given cluster. Loops were chosen from well-populated clusters. LoopAB, loopBC, and loopCD are two residues and constrained in the residue file to a Gly-Asp ($\alpha_L\beta$) motif. This ensured favorable N-capping and C-capping interactions to support the folded core.[6-7]

**COMBS and vdM sampling.** All COMBS code can be found here: https://github.com/npolizzi/Combs

**Flexible Backbone Sequence Design.** We wrote a RosettaScript for flexible backbone sequence design, implemented in Rosetta 3.5, that progresses through a cycle of backbone and side-chain relaxation and fixed backbone sequence design, with a filtering step based on packing score (see RosettaScript below).

***Amino acids allowed to vary during design.*** To avoid deleterious oxidations within the active site we disallowed Cys and Met residues throughout the protein. Additionally, no His residues were allowed. The residue files (resfile_ex1.txt, resfile_ex3.txt, resfile_aro.txt) are provided below showing exactly which residues were allowed to be designed. The binding site H-bonding residues Asn16, Asn75, Gly9, and Gly68 and second-shell Asn45 and Asn104 were held constant. In addition, the loop motifs (Gly/Asp) were held constants for all three loops. Rosetta was able to place two Trp residues in close proximity to the NDI (Trp 13 and Trp 20) and they were then constrained in the resfile.

From the NBP-1 sequence, oxidizable aromatic residues near the NDI were varied for two additional designs: for NBP-2 Trp 13 was replaced with a Tyr and Trp 20 was unchanged. For NBP-3 Trp13 and Trp20 were kept, and a buried Tyr (position 79) was constrained.

***Flexible backbone design protocol*.** Distance constraints (constraints.cst) between the H-bonding Asn and NDI-carbonyl oxygens were loaded, the sequence was designed with a fixed backbone, the sidechains and backbone were minimized, three trials of a Monte Carlo flexible backbone subprotocol were performed followed by one trial of an additional Monte Carlo flexible backbone protocol with additional rotamers allowed, and 100 models with packing score (pstat) > 0.48 were output. Designs were visualized with PyMol and analyzed for total score and pstat to choose NBP-1.

***Flexible backbone design sub-protocol*.** The flexible backbone design sub-protocol (see below, rscript_flexbb_relax.xml) consists of three Monte Carlo trials of (1) fixed backbone design with ref2015 weights and the addition of extra rotamer sampling around $\chi_1$ (ex1, level 1; i.e. sampled between 1 standard deviation of the mean chi angle value for reach rotamer) and $\chi_2$ (ex2, level 1) sidechain dihedrals, (2) fast relax minimization (with increased repulsive features from KillA2019 modifications). At the end of step (2), the model is filtered for native-like structure packing with PackStat (if 1 of 3 trials the PackStat score is > 0.48 the model passes the filter). This model then undergoes an additional Monte Carlo trial of (1) fixed backbone design as above, but with rotamers sampled between 2 full standard deviations of the mean chi angle (ex1, level 3) and sidechain dihedrals (ex2, level 3), (2) rosettacon2018-modified FastRelax, and (3) a final PackStat filter required pstat > 0.48. The final designed sequences of NBP-1, NBP-2, and NBP-3 selected for expression were as follows:

>NBP-1
SAKQDFAEGVKLWQENAVLWTRLVQAFQSGDQSTVDSLLKQLDANAARVEQLLQRIIS
ETGDELARKGESLFQRNQQLFSQLKTLFSQGDEDTAKAVLEEIQSNLNQIQQIITEAQKR
L

>NBP-2
SSAKSDFTEGVVLWQQNATLWNKLVQAFQKGDQSTVQSLLKQLDANAARVEQLLQRII
SETGDELARRGEELFQRNQQLYSQLKQLFSEGDEDTAKQVLEEIQANLSQIQAIISTAQK
RL

>NBP-3
SSAKQQFSEGVALYQTNTVLWQKLVVAFQQGDESTVSSLLKQLDANANRVEQLLQQIIS
ETGDELARRGEQLFQRNQELFSQLKELFTQGDRDTAKQVLEEIQANLAQISVIISQAAKK
L

**Ab initio folding.** Rosetta *ab initio* folding prediction calculations[8] were performed on the NBP-1 sequence in Rosetta 3.5. 10,000 structures were generated. Cα RMSD of the helical backbone was scored against residues 6-28, 33-59, 64-86, and 92-115 of the design model (Fig. S4).

**Chai-1 Structure Prediction.**[9] The Chai-1 webserver was used to predict the structure of NDI1-NBP1. This tools utilizes a protein sequence and SMILES string of the ligand to predict the protein-ligand structure. NDI1-NBP1 was very well predicted with this tool giving backbone RMSD of 1.07 Å and buried side-chain RMSD of 1.45Å with a confidence score (interface predicted template modeling (ipTM) score) of 0.88. It also placed the ligand precisely where designed. We also used this tool to predict the NDI2-NBP1 structure (which was not originally designed) and found that it, too, matched very closely to the NDI1-NBP1 structure. It gave a backbone RMSD of 0.96 Å and buried sidechain rmsd of 1.36 Å with an ipTM score of 0.86.

**Size exclusion chromatography.** Gel filtration profiles were obtained using a Superdex 75 5/150 column on an FPLC system (GE Healthcare AKTA). 500 µL of 100 µM apo-NBP-1 or 200 µM holo-MPP1 was injected onto the column and eluted with a 50 mM NaPhos, 150 mM NaCl, pH 7.5 buffer mobile phase at a flow rate of 0.2 mL/min (Fig. S13).

**Circular dichroism (CD).** CD spectra were collected on a Jasco J-810 CD spectrometer in a 0.1 cm path length quartz cuvette. Spectra were collected over a wavelength range from 200 to 450 nm. Apo- and holo-NBP-1 were prepared at 8 µM for 200-250 nm window and 500 µM for the visible region in 50 mM NaPhos, 150 mM NaCl pH 7.5.

**Electronic absorption spectroscopy.** Electronic absorption spectra were collected using an HP 8453 UV-Vis spectrophotometer or Cary 300 Bio spectrophotometer in a low-volume (750 μL) 1-cm path-length quartz cuvette.

**Visualization of protein structures and image rendering.** Protein models were visualized and rendered in the PyMol visualization program.[10]

**Protein expression and purification.** The plasmid coding for the protein sequence of NBP-1 was ordered from GenScript, which was cloned into the IPTG-inducible pET-11a plasmid (cloning site NdeI-BamHI). The sequence also coded for an N-terminal 6xHis-tag followed by a TEV protease cleavage sequence, followed finally by the designed sequence. The cloned gene sequence is:
ATGCATCACCACCACCACCATGAAAATCTATACTTCCAATCGTCTGCGAAGCAAGAT
TTTGCCGAAGGTGTTAAACTGTGGCAGGAGAACGCGGTCCTGTGGACCCGTTTGGTT
CAGGCGTTTCAATCTGGCGACCAGTCCACCGTTGATAGCCTGCTGAAGCAGCTGGAC
GCTAATGCCGCGCGCGTGGAACAGTTGTTACAGCGCATTATTAGCGAAACTGGTGAC
GAGCTCGCTCGTAAAGGCGAGAGCTTGTTTCAGCGTAATCAGCAGCTGTTCAGCCAG
CTTAAGACGCTGTTCAGCCAAGGTGACGAGGATACCGCAAAAGCGGTGTTGGAAGA
GATCCAATCCAACCTGAACCAAATTCAACAAATCATCACCGAAGCACAGAAACGTC
TG

The expressed protein sequence was finally:
MHHHHHHENLYFQS/SAKQDFAEGVKLWQENAVLWTRLVQAFQSGDQSTVDSLLKQL
DANAARVEQLLQRIISETGDELARKGESLFQRNQQLFSQLKTLFSQGDEDTAKAVLEEIQ
SNLNQIQQIITEAQKRL

where the "/" defines the cleavage site of TEV protease. The plasmids were transfected into *E. coli* BL21(DE3) cells, which were grown in LB/carbenicillin media until OD @ 600 nm = 0.6. The cells were then induced with 1 mM IPTG (final concentration) and allowed to grow for 4 more hours. Cells were then centrifuged and frozen. The frozen cell pellets were lysed via sonication in 50 mM NaPhos, 150 mM NaCl, 20 mM imidazole pH 7.5. The expressed, His-tagged NBP-1 protein was purified via a Ni NTA column and confirmed by gel electrophoresis, with an approximate yield of 20 mg/L. NBP-2 and NBP-3 were expressed and purified in the same manner.

For NMR experiments, cells were grown in M9 minimal media with isotope-labeled ammonia ($^{15}N$) and glucose ($^{13}C$) from Cambridge Isotopes until OD at 600 nm = 0.6 before centrifugation and resuspension and induction in LB media as described above.

**NMR spectroscopy.** All NMR samples contained 0.2 mM protein in a buffer (50 mM NaPhos, 150 mM NaCl, pH 7.5) and 5% $D_2O$. 1D Spectra were recorded at 298 K on a Bruker 800 MHz spectrometer equipped with a cryogenic probe. The $^{1}H$ zgesgp spectra were recorded with 512 scans with 30 ppm spectral width. $^{1}H$ chemical shifts were referenced with respect to the residual water peak at 4.75 ppm. All spectra were processed and analyzed using the program TopSpin 3.6

(Bruker, Karlsruhe, Germany). Prior to Fourier transformation, time domain data were multiplied by sine square bell window functions shifted by 90º and zero-filled once.

Sequence specific backbone ($^{1}$HN, $^{15}$N, $^{1}$Hα, $^{13}$Cα) and $^{1}$Hb / $^{13}C_{\beta}$ resonance assignments were obtained the program ARTINA. Assignments were obtained for 92.3% of the backbone (excluding the N-terminal $NH_3$+, the Pro $^{15}$N and the $^{13}$C' preceding prolyl residues) and $^{13}C_{\beta}$, and for 69.3% of the side chain chemical shifts (excluding Lys NH3+, Arg $NH_2$, OH, side chain $^{13}$C' and aromatic $^{13}$Cγ) which are assignable with the set of NMR experiments provided above. Chemical shift table has been deposited into BMRB (BMRB ID is 52600).

Figure 6A displays a single peak for H11, H12, H13, and H14 due to chemical shift degeneracy, a consequence of the structural symmetry of the ligand. In contrast, Figure 6B reveals that H11, H12, H13, and H14 exhibit distinct chemical shifts, indicating different chemical environments when the ligand is in the presence of the designed protein. Figure 10A was recorded using the pulse sequence mlevphpp for homonuclear Hartmann-Hahn transfer, employing the MLEV17 sequence with an 80 ms mixing time. Figure 10B, on the other hand, was recorded using the pulse sequence dipsi2gpphwgxf for 2D homonuclear Hartmann-Hahn transfer, utilizing the DIPSI2 sequence with an 60 ms mixing time in the presence of a $^{13}$C and $^{15}$N filter in both the F1 and F2 dimensions. This filtering technique (H[$^{12}$C, $^{14}$N] (t1) -- H[$^{12}$C, $^{14}$N] (t2)) effectively removes all signals from the $^{13}$C, $^{15}$N-labeled protein, thereby allowing the observation of the distinct chemical shifts of the non-labeled ligand's protons.

**Molecular Dynamics (MD) and Simulations Parameters.** Each NDI-binder design was simulated to assess stability of the complex. The starting structures were derived from the Rosetta models, and protein residues were assigned their standard protonation states at pH 7. $Ph_2$-NDI (NDI-1) was parameterized using Gaussian09 and the Antechamber program within Amber22.[11-13] The partial charges of the molecule were determined by first optimizing its geometry at the B3LYP/6-31G* level, followed by calculating the electrostatic potential using the Merz-Singh-Kollman method in Gaussian09.[11] Charge fitting was then performed using Antechamber's RESP program within AmberTools.[14-15] All other small molecule parameters were assigned by Antechamber based on the GAFF2 database.[16-17]

The MD system was prepared using Amber's tleap program. Protein residues were parameterized using the ff19SB forcefield[18], and each complex was solvated with OPC model waters[19] in a box with 10Å padding from the protein. Sodium and chloride ions were added to reach a charge-neutral 150 mM NaCl concentration to match experimental conditions. All simulations were performed in Amber22 and began with 1,000 restrained steepest-descent minimization steps, followed by a maximum of 7,000 steps of conjugate gradient minimization. The system was then heated up from 150K to 293 K over 50 ps in the NVT ensemble with the Langevin thermostat and a 1 fs integration timestep. The system was then switched to the NPT ensemble, and pressure was maintained at 1 atm using the Monte Carlo barostat. The protein and ligand were initially restrained with harmonic potential at a 10 kcal/(mol·Å$^2$) force constant, which was gradually reduced to 0 kcal/(mol·Å$^2$) over 8 equilibration steps, totaling 1ns.

Following minimization and equilibration, an unrestrained production run was conducted for 1 microsecond under periodic boundary conditions with a 2 fs integration timestep. The

SHAKE algorithm[20] was used to restrain hydrogens, the Particle Mesh Ewald method[21] was used to calculate long-range electrostatics, and non-bonded electrostatic and Lennard-Jones interactions were cut off at 10 Å. Three independent simulations were performed for each design.

**Pump-probe time-resolved transient optical**. Ultrafast transient absorption spectra were obtained through standard pump-probe method. Optical pulses (≥ 80 fs, 1 kHz) centered at 800 nm were generated with Solstice Ace (Spectra-Physics, Milpitas, CA, USA), which consisted of a regenerative amplifier seeded by a mode-locked Ti:Sapphire oscillator. A fraction of the output from the regenerative amplifier was split to feed an optical parametric amplifier TOPAS-C (Light Conversion, Vilnius, Lithuania), which generates pump pulses whose center wavelength was tuned for each measurement. The pump beam was fed into HARPIA spectroscopy system (Light Conversion, Vilnius, Lithuania). HARPIA chopper was set at *f*/4 (250 Hz). Another fraction of the regenerative amplifier output was fed into HARPIA, passed through an 8-ns optical delay line, focused onto a 3 mm calcium fluoride window for supercontinuum generation, and used as probe beam. The polarization and attenuation of the pump and probe beams were controlled by half-wave plate and Glan-Taylor prism pairs. Between pump and probe beams, the polarization was set for the magic angle (54.7°). The pump beam was focused onto the sample cuvette with an f = +100 cm lens, while the probe beam was focused with a concave mirror. The spot size diameter was ~0.2 – 0.3 mm. The pump power utilized was about 900 µW. The probe beam after passing through the sample was adjusted with an ND filter to avoid detector saturation. Then the probe beam was focused onto the entrance slit of spectrograph Kymera 328i (Andor Technology, Belfast, UK), which was interfaced by a broadband Si n-channel metal-oxide semiconductor (NOMS) detector (S3901 + C7884, Hamamatsu Photonics, Hamamatsu City, Shizuoka, Japan). All measurements utilized a 5 mm path length quartz sample cell. Samples in cuvette were purged under nitrogen for above 30 minutes. All transient spectra reported represent averages obtained over 5 scans, with each scan consisting of over 200 time delays. Following all pump-probe transient absorption experiments, electronic absorption spectroscopy was utilized to verify the compound integrity. Data were processed with Surface Xplorer (Ultrafast System, Sarasota, FL, USA), and global fitting analysis was performed using CarpetView (Light Conversion, Vilnius, Lithuania).

**Infrared (IR) Spectroscopy.** IR spectra were recorded on a Thermo Scientific Nicolet iS50 FTIR Spectrometer. 6 mM samples were prepared of NDI1 in DCM, apo-NBP1 (15N-13C labeled), and NDI1-NBP1 (15N-13C labeled) in 50 mM NaPhosphate 150 mM NaCl pH 7.5 buffer. Measurements were made in a room temperature OTTLE cell[22] consisting of $CaF_2$ windows, 50 µm spacer, Au mesh working electrode, Ag mesh reference electrode, and Pt wire counter electrode.

**Determination of apparent dissociation constant.** We used spectral titration experiments to determine the binding dissociation constants for NDI**1** in NBP1. Initial experiments were performed using UV-vis to monitor the NDI absorbance at 382 nm. Due to the low solubility of NDI1 in buffer, a 3µM NDI1 solution was prepared in 10% DMSO:buffer (buffer = 50 mM NaPhos 150 mM NaCl pH 7.5). NBP1 was serially titrated in from a 131 µM stock solution and showed binding in a 1:1 stoichiometry (Fig. 3B).

In addition, we performed tryptophan fluorescence quenching experiments[23] to determine apparent $K_d$ of NDI1-NBP1. We used a BioTek Synergy 2 Neo2 plate reader with excitation wavelength of 295 nm and emission wavelength of 355 nm. Aliquots of NBP1 (98 μM) solution were added to NDI1 solutions at 245 nM, 490 nM, 760 nM, and 980 nM. For NBP1-AA (194 μM), aliquots were added to solution of NDI1 at 485 nM and 730 nM. The apparent $K_d$ for each protein was determined by globally fitting the change in emission at 355 nm.

**Photolabeling Experiments.**

***General methods and instrumentation*:** Illumination was performed using a Penn PhD Photoreactor M2 (Sigma Aldrich, Z744035) with a 450 nm blue light source module (Sigma Aldrich, Z744033). Fan speed was set at 6800 rpm under manual control with 100/min stirring and samples were illuminated at 100% intensity for indicated time. Bovine serum albumin (BSA) was purchased from Rockland (BSA-10). Labeling probes were prepared as previously reported.[24]

***Recombinant protein biotinylation assay*:** In order to test both self- and intermolecular protein labeling, biotinylation experiments were performed with NDI binders alone or with BSA. A 25 μL reaction system in PBS was prepared with 10 μM binder or NDI compound with or without BSA. Labeling probes (diazirine-biotin, aryl-azide-biotin, biocytin-hydrazide, or biotin-phenol) was then added into the solution to reach a final concentration of 100 μM and mixed thoroughly before illumination with LED for indicated time at 4 °C. Afterwards, sample loading buffer was added, and the samples were subjected to SDS-PAGE.

***Western blot protocol*:** For immunoblotting analysis, proteins were loaded on 4-12% BisTris gels (Bolt 4-12% 17-well, Thermo Fischer, NW04127BOX), and transferred from SDS-PAGE gels to PVDF membranes (Thermo Fischer, IB24002) using an iBlot-2 dry blotting system (Thermo Scientific, IB21001). For total protein visualization, Ponceau S staining (Tocris, 5225) was used. Membranes were then washed with Tris buffered saline (TBST, 37mM sodium chloride, 20mM Tris, 2.7mM potassium chloride, 0.05% Tween 20; pH=7.4) and blocked with TBST containing 0.1% Tween-20 and 5% BSA. In order to visualize protein biotinylation, LIC-OR IRDye 800CW streptavidin antibody (LI-COR Biosciences, 926-32230) was added. Immunoblots images were captured by an infrared LI-COR imager (LI-COR Biosciences, Odyssey CLx). In-gel fluorescence and immunoblot fluorescence signals were detected on a BioRad imager (ChemiDoc XRS+ System). Data were analyzed using GraphPad Prism (v8.0.1), ImageStudioLite (v5.2.5), and Adobe Illustrator (v22.1).

**Preparation of NDI derivatives.**

***$Ph_2$-NDI (NDI1).*** NDI1 was purchased from Sigma and used without further purification.

***Synthesis of $(C_5H_{11})_2$-NDI (NDI2*).** A suspension of 1,4,5,8-Naphthalenetetracarboxylic dianhydride (0.500 g, 1.32 mmol) and 3-aminopentane (1.15 g, 13.2 mmol) in dimethylformamide (15 mL) was heated to 120 °C and reacted for 24 hours. The reaction was cooled to room temperature precipitating crude 8. The precipitate was filtered and washed with MeOH (3 x 10 mL) and diethyl ether (3 x 10 mL) yielding pure NDIAP. Yield: 443 mg (83%). MALDI-TOF: obsd, 405.319 $(M-H)^-$; calcd, 406.48 ($M = C_{24}H_{46}N_2O_4$); $^1H$ NMR (500 MHz,

$CDCl_3$): *d* 8.67 (s, 4H), 4.93-4.87 (m, 2H), 2.18-2.09 (m, 4H), 1.90-1.81 (m, 4H), 0.83-0.80 (t, 12H)).

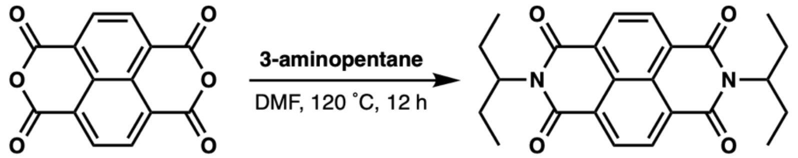


**Scheme S1.** Synthesis of $(C_5H_{11})_2$NDI (NDI2).

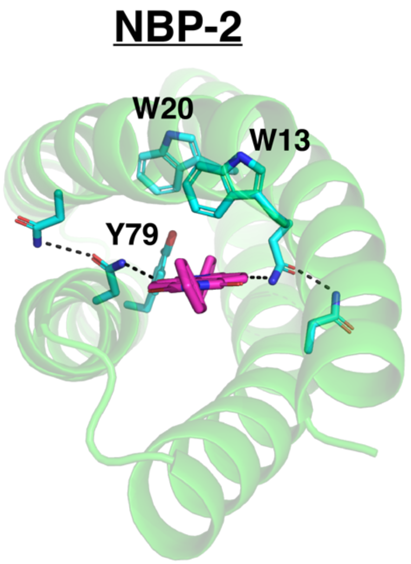


**Figure S1.** Design models of NBP-2 and NBP-3 showing the different aromatic residues near the NDI cofactor (pink).

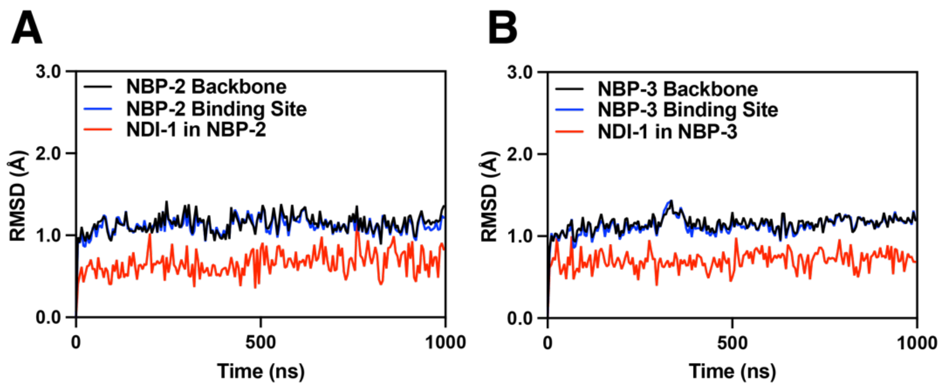


**Figure S2.** Averaged RMSD of NDI1-NBP2 (A) and NDI1-NBP3 (B) complexes over three 1 µs simulations showing extreme stability of the design and impressive RMSD for NDI1 cofactor within the binding site. Binding site is backbone within 10 Å of the cofactor.

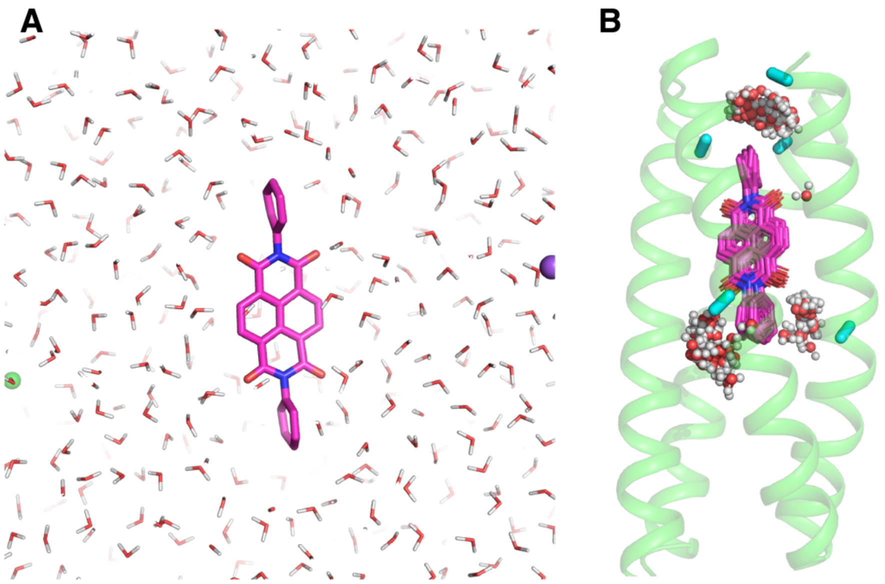


**Figure S3.** Comparison of hydration sphere around NDI in NBP-1 from MD simulations. (A) Shows waters within 15 Å of "free" NDI-1 in aqueous buffer (green spheres are chloride anions and purple spheres are sodium cations) showing water molecules surrounding the NDI. (B) Shows the clusters of water that persist around the NDI over the 1µs simulation. 30 ns frames were clustered on the protein backbone (cartoon showed is the centroid) revealing three water clusters (near small Ala residues in cyan), illustrating how dry the binding site is relative to free NDI-1.

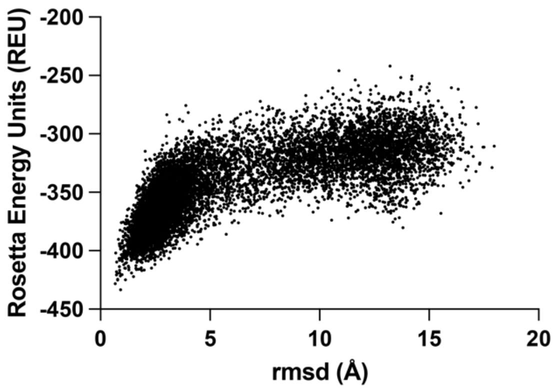

**Figure S4.** Rosetta *ab initio* folding predictions for NBP-1.

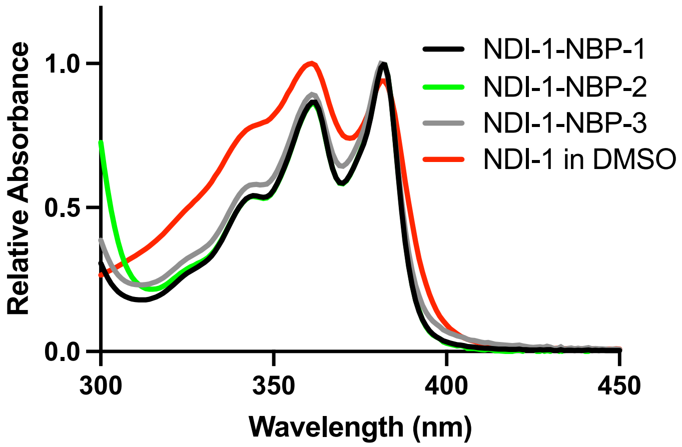

**Figure S5.** Electronic absorption spectra of NDI-1 bound to NBP1-3 showing similar absorption changes relative to NDI-1 in DMSO.

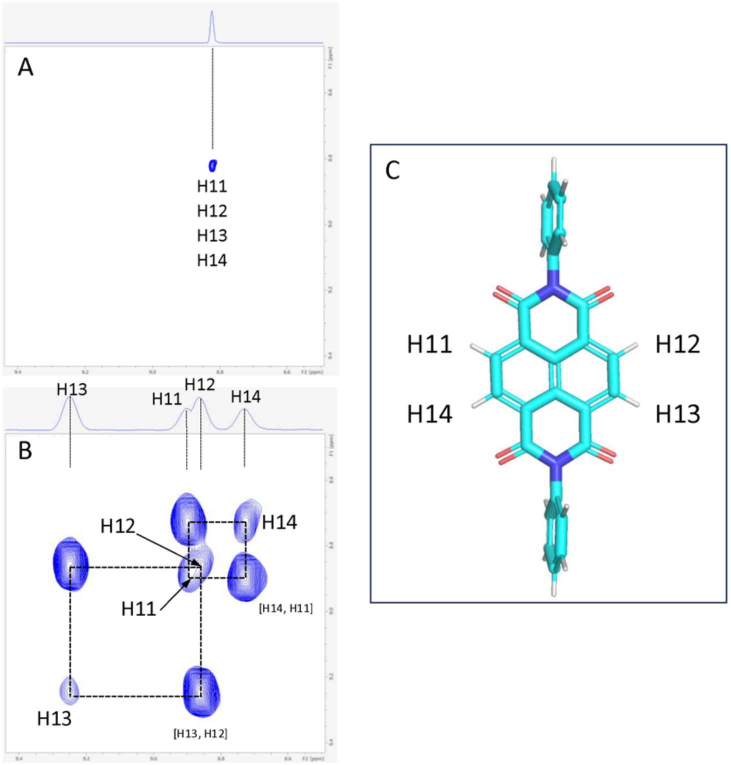


**Figure S6.** 2D $^{1}$H NMR spectrum at 800 MHz of NDI1 showing the naphthalene protons go from a single peak outside of the protein to multiple within the protein binding site. (A) A section of the $^{1}$H-TOCSY spectrum of free NDI-1 (in DMSO). (B) 2D homonuclear X-filtered TOCSY (using the pulse sequence dipsi2gpphwgxf) for the non-labeled ligand in the presence of the $^{13}$C, $^{15}$N-labeled NBP1. (C) Chemical structure of ND1 with relevant naphthalene protons labeled.

Table S1. Ring Current Effects on Chemical Shifts for Residues Near NDI

| Residue | Atom | Observed Shift (PPM) | BMRB average shift (PPM) | Difference/Standard Deviation |
|---|---|---|---|---|
| ILE 112 | HD1 | -1.298 | 0.683 ± 0.279 | 7.10 |
| ILE 108 | HG2 | -0.832 | 0.778 ± 0.266 | 6.05 |
| LEU 53 | HD1 | -0.331 | 0.755 ± 0.272 | 3.99 |
| VAL 49 | HG2 | -0.739 | 0.806 ± 0.276 | 5.60 |

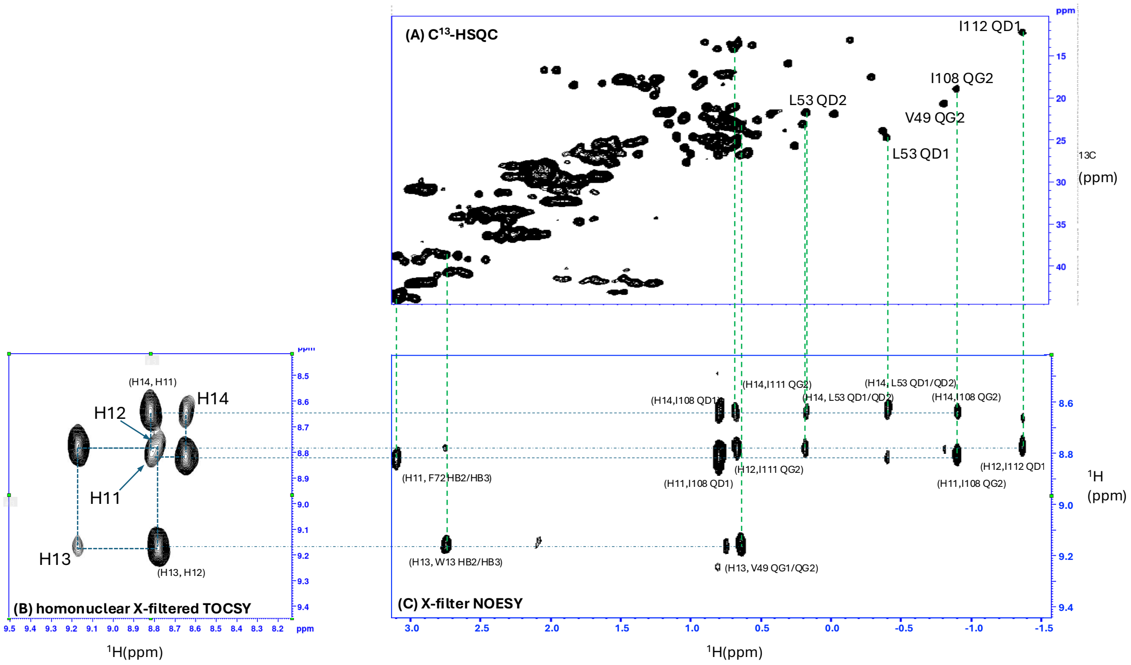


**Figure S7.** The strong ring current of the naphthalene core of NDI1 results in upfield shifted signals from residues within the NDI binding site. (**A**) $^{1}H$-$^{13}C$ HSQC spectrum of NDI1-NBP1. (B) Homonuclear X-filtered TOCSY aligned with (C) X-filtered NOESY showing degenerate signals for the naphthalene protons interacting with residues close to the binding site and shifting their signals upfield.

Table S2. Intermolecular NOE Distances Between NDI1 and NBP1

| NDI1 Atom | NBP1 Atom | Distance (Å) |
|---|---|---|
| H14 | I108 HD1 | 5 |
| H14 | I111 HG2 | 5 |
| H14 | L53 HD1/L53HD2 | 6 |
| H14 | L53 HD1/L53HD2 | 6 |
| H14 | I108 HG2 | 5 |
| H11 | F72 HB2 | 6 |
| H11 | F72 HB3 | 6 |
| H11 | I108 HD1 | 5 |
| H12 | I111 HG2 | 5 |
| H11 | I108 HG2 | 5 |
| H12 | I112 HD1 | 5 |
| H13 | W13 HB2 | 6 |
| H13 | W13 HB3 | 6 |
| H13 | V49 HG1 | 6 |
| H13 | V49 HG2 | 6 |
| H1 | A46 HB | 5 |
| H3 | A46 HB | 5 |
| H4 | A115 HB | 5 |
| H6 | A115 HB | 5 |

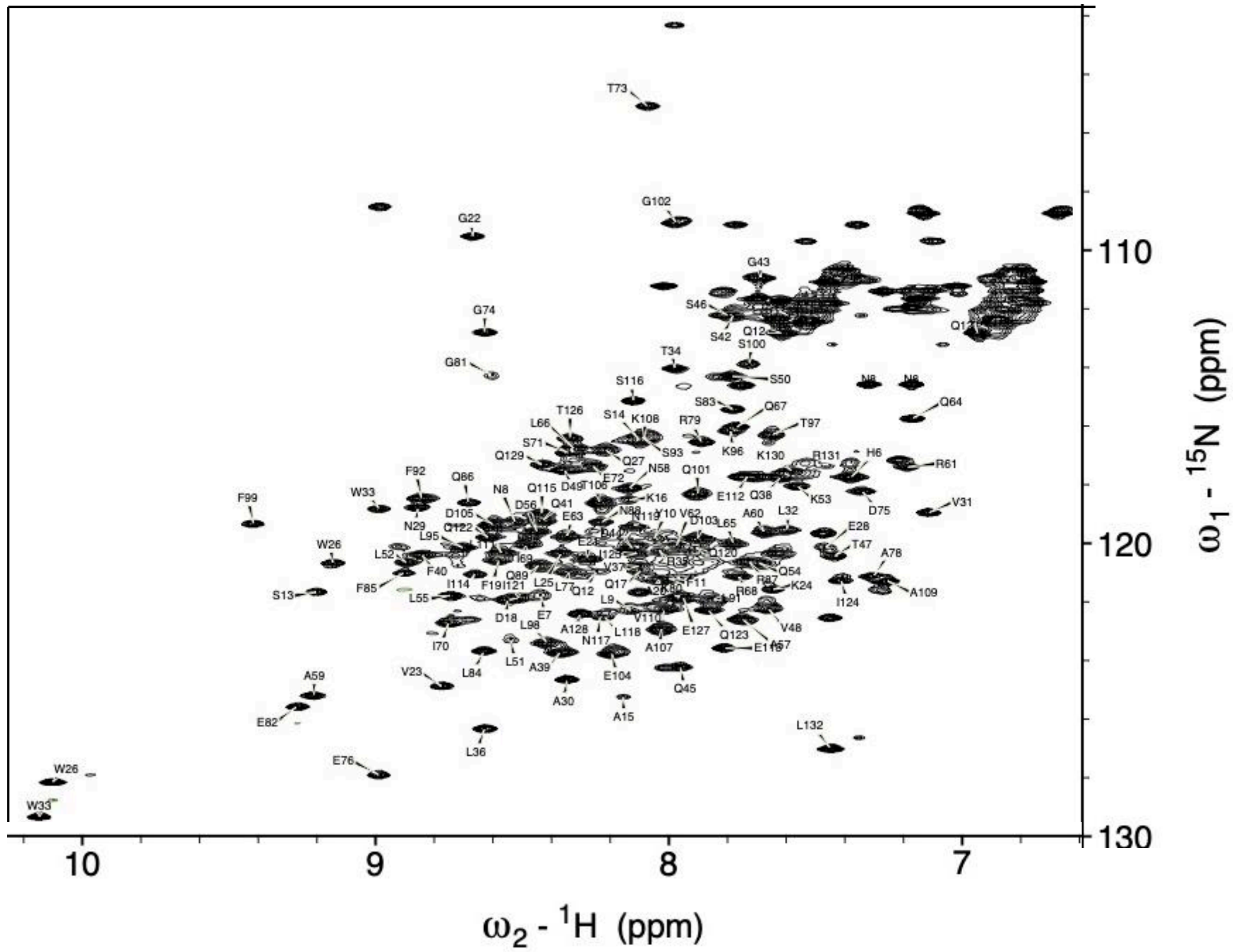


**Figure S8.** Well-resolved 1H-15N HSQC spectrum at 800 MHz of NDI1-NBP1 showing assigned signals.

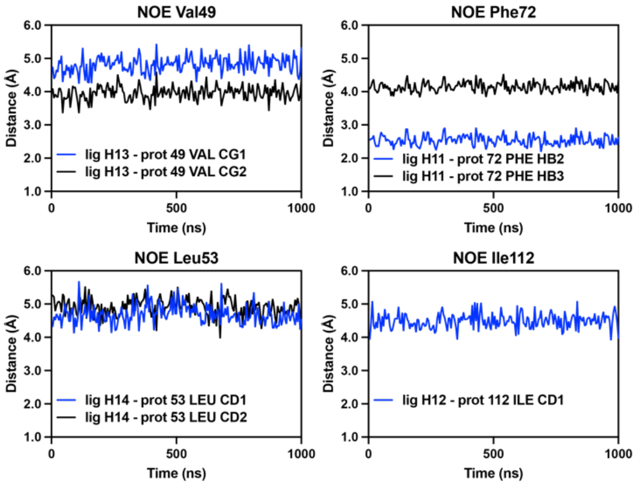


**Figure S9.** Analysis of MD simulations showing atom-atom distances over the course of MD trajectory corroborating NOEs calculated from NMR.

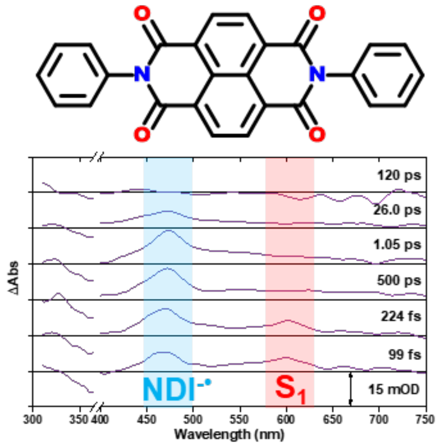


**Figure S10.** Pump-probe transient dynamics of $NDIPh_2$ displaying intramolecular charge separated species $NDI^{-\bullet}$ (blue) forming as the S1 excited state absorption (red) decays. Experimental conditions: Dichloromethane; ambient temperature; $\lambda_{ex}$= 380 nm

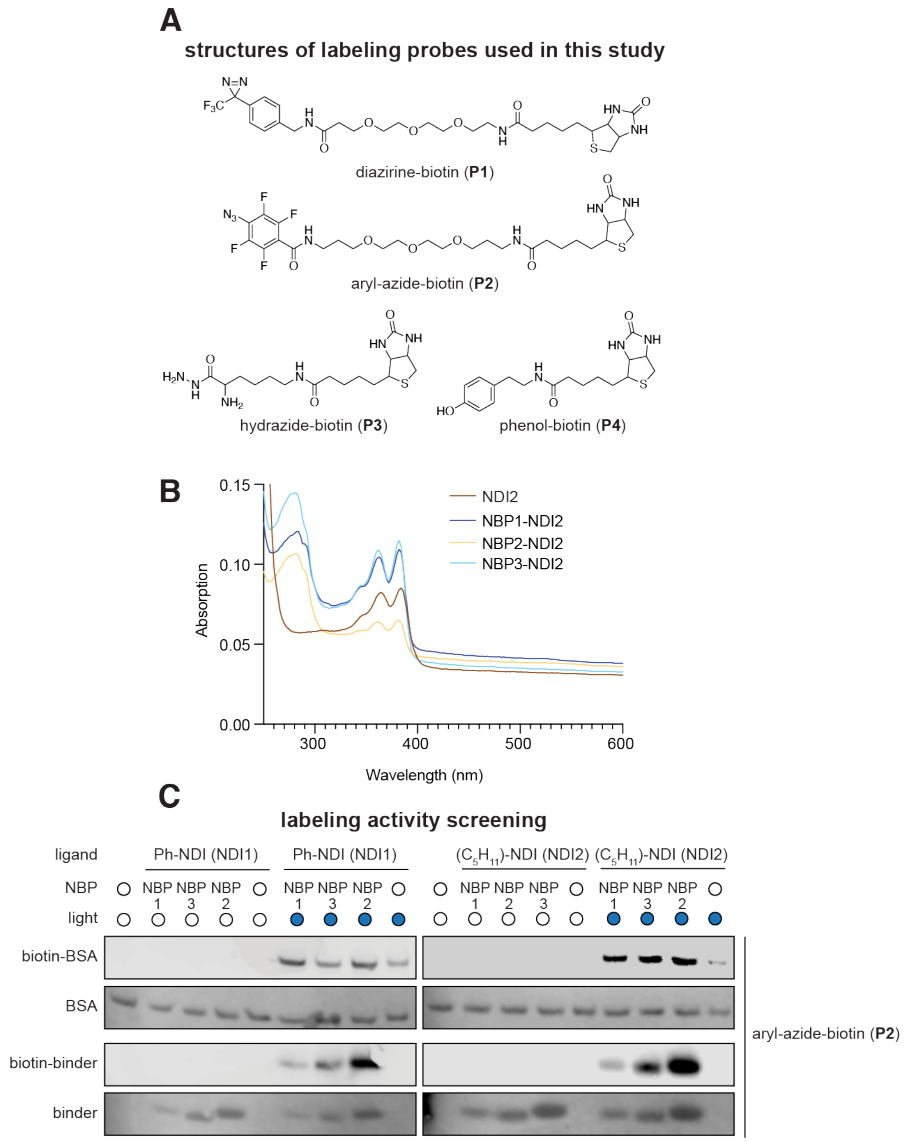


**Figure S11.** (A) Structures of labeling probes used in this study. (B) Electronic absorption spectra showing after irradiation NDI binders retain their absorption spectra. (C) Labeling activity screen with both NDI derivatives and all three designs. They all showed the ability to activate aryl-azide-biotin (P2) to self-label as well as label BSA.

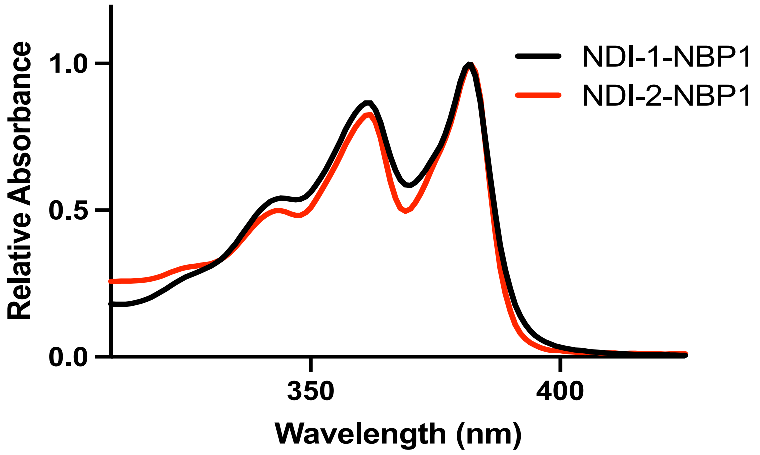


**Figure S12.** Electronic absorption spectra showing similar absorbance features for the Ph2NDI (NDI-1) and alkyl-NDI (NDI-2) when bound to NBP1.

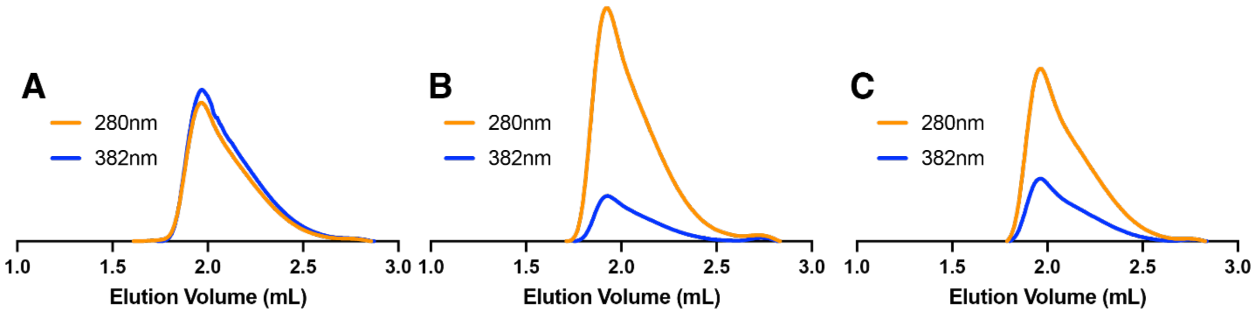


**Figure S13.** FPLC SEC chromatograms of (A) NBP1, (B) NBP2, and (C) NBP3.

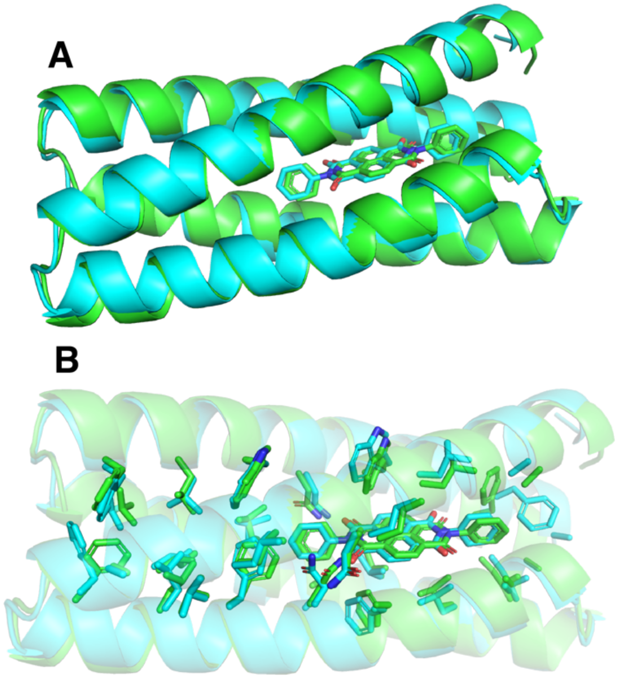


**Figure S14.** Overlays of Chai-1 predicted ligand-protein complexes. (A-B) Chai-1 prediction (teal) and NDI1-NBP1 design model (green) showing close match to the backbone (A) and buried sidechains (B). (C-D) Chai-1 prediction of NDI2 bound to NBP1 (blue) and NDI1-NBP1 design model (green) showing close match to the backbone (A) and buried sidechains (B).

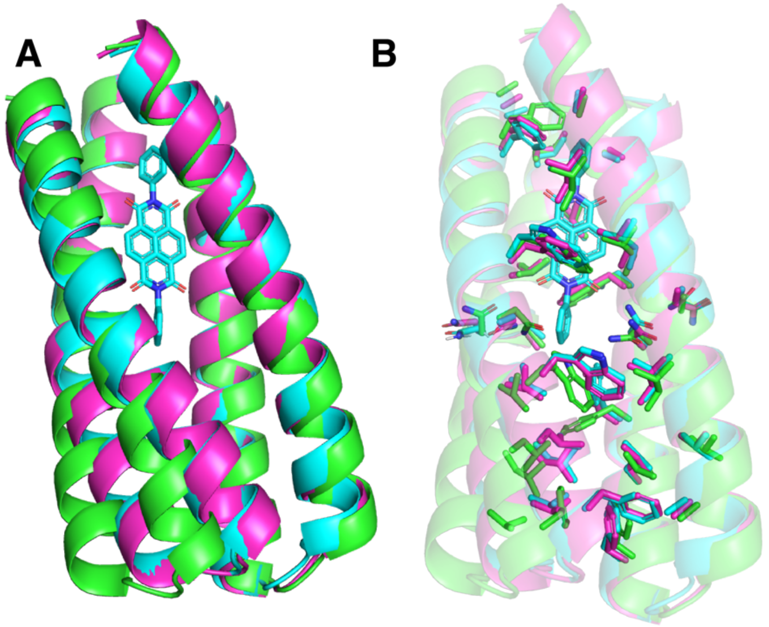


**Figure S15.** Overlay of NDI1-NBP1 design model with different structure prediction software. (A) Backbone comparison and (B) buried sidechain comparison of ND1-NBP1 design model (teal), AlpaFold2[25] model (purple), and ESMfold[26] prediction (green).

**Command lines, RosettaScript, and flags for flexible backbone sequence design protocol.**

**Resfiles**

**resfile_aro.txt**
NATRO
ALLAAxc
NOTAA MTC
USE_INPUT_SC
start

16 A PIKAA N EX ARO 1 LEVEL 3 EX ARO 2 LEVEL 3
75 A PIKAA N EX ARO 1 LEVEL 3 EX ARO 2 LEVEL 3

30 A PIKAA G EX ARO 1 LEVEL 3 EX ARO 2 LEVEL 3
31 A PIKAA D EX ARO 1 LEVEL 3 EX ARO 2 LEVEL 3

61 A PIKAA G EX ARO 1 LEVEL 3 EX ARO 2 LEVEL 3
62 A PIKAA D EX ARO 1 LEVEL 3 EX ARO 2 LEVEL 3

89 A PIKAA G EX ARO 1 LEVEL 3 EX ARO 2 LEVEL 3
90 A PIKAA D EX ARO 1 LEVEL 3 EX ARO 2 LEVEL 3

1 A PIKAA S EX ARO 1 LEVEL 3 EX ARO 2 LEVEL 3
2 A PIKAA NVAEKTSDRQ EX ARO 1 LEVEL 3 EX ARO 2 LEVEL 3
3 A PIKAA NVATSQRK EX ARO 1 LEVEL 3 EX ARO 2 LEVEL 3
4 A PIKAA NVATSQ EX ARO 1 LEVEL 3 EX ARO 2 LEVEL 3
5 A PIKAA NVAETSDQ EX ARO 1 LEVEL 3 EX ARO 2 LEVEL 3
6 A PIKAA FVAWITLYGS EX ARO 1 LEVEL 3 EX ARO 2 LEVEL 3
7 A PIKAA NVATSQ EX ARO 1 LEVEL 3 EX ARO 2 LEVEL 3
8 A PIKAA NVAETSDQ EX ARO 1 LEVEL 3 EX ARO 2 LEVEL 3
9 A PIKAA G EX ARO 1 LEVEL 3 EX ARO 2 LEVEL 3
10 A PIKAA NVAKTSRQ EX ARO 1 LEVEL 3 EX ARO 2 LEVEL 3
11 A PIKAA NVAKTSRQ EX ARO 1 LEVEL 3 EX ARO 2 LEVEL 3
12 A PIKAA FVAWITLYGSNQ EX ARO 1 LEVEL 3 EX ARO 2 LEVEL 3
13 A PIKAA W EX ARO 1 LEVEL 3 EX ARO 2 LEVEL 3
14 A PIKAA NVATSQ EX ARO 1 LEVEL 3 EX ARO 2 LEVEL 3
15 A PIKAA NVAETSDQ EX ARO 1 LEVEL 3 EX ARO 2 LEVEL 3
17 A PIKAA NVAETSDQ EX ARO 1 LEVEL 3 EX ARO 2 LEVEL 3
18 A PIKAA NVAKTSRQ EX ARO 1 LEVEL 3 EX ARO 2 LEVEL 3
19 A PIKAA NVFAWEIKTLYSDRQ EX ARO 1 LEVEL 3 EX ARO 2 LEVEL 3
20 A PIKAA W EX ARO 1 LEVEL 3 EX ARO 2 LEVEL 3
21 A PIKAA NVATSQ EX ARO 1 LEVEL 3 EX ARO 2 LEVEL 3
22 A PIKAA NVAKTSRQ EX ARO 1 LEVEL 3 EX ARO 2 LEVEL 3

23 A PIKAA FVAWITLYGS EX ARO 1 LEVEL 3 EX ARO 2 LEVEL 3
24 A PIKAA NVAETSDQ EX ARO 1 LEVEL 3 EX ARO 2 LEVEL 3
25 A PIKAA NVATSQ EX ARO 1 LEVEL 3 EX ARO 2 LEVEL 3
26 A PIKAA NVAKTSRQ EX ARO 1 LEVEL 3 EX ARO 2 LEVEL 3
27 A PIKAA FVAWITLYGS EX ARO 1 LEVEL 3 EX ARO 2 LEVEL 3
28 A PIKAA NVAETSDQ EX ARO 1 LEVEL 3 EX ARO 2 LEVEL 3
29 A PIKAA ST EX ARO 1 LEVEL 3 EX ARO 2 LEVEL 3
31 A PIKAA NVAEKTGSDRQ EX ARO 1 LEVEL 3 EX ARO 2 LEVEL 3
32 A PIKAA NVAEKTSDRQ EX ARO 1 LEVEL 3 EX ARO 2 LEVEL 3
33 A PIKAA NVATSQ EX ARO 1 LEVEL 3 EX ARO 2 LEVEL 3
34 A PIKAA NVAETSDQ EX ARO 1 LEVEL 3 EX ARO 2 LEVEL 3
35 A PIKAA FVAWITLYGS EX ARO 1 LEVEL 3 EX ARO 2 LEVEL 3
36 A PIKAA NVAETSDQ EX ARO 1 LEVEL 3 EX ARO 2 LEVEL 3
37 A PIKAA NVATSQ EX ARO 1 LEVEL 3 EX ARO 2 LEVEL 3
38 A PIKAA FVAWITLYGS EX ARO 1 LEVEL 3 EX ARO 2 LEVEL 3
39 A PIKAA FVAWITLYGS EX ARO 1 LEVEL 3 EX ARO 2 LEVEL 3
40 A PIKAA NVAKTSRQ EX ARO 1 LEVEL 3 EX ARO 2 LEVEL 3
41 A PIKAA NVAKTSRQ EX ARO 1 LEVEL 3 EX ARO 2 LEVEL 3
42 A PIKAA FVAWITLYGS EX ARO 1 LEVEL 3 EX ARO 2 LEVEL 3
43 A PIKAA NVAETSDQ EX ARO 1 LEVEL 3 EX ARO 2 LEVEL 3
44 A PIKAA NVATSQ EX ARO 1 LEVEL 3 EX ARO 2 LEVEL 3
45 A PIKAA N EX ARO 1 LEVEL 3 EX ARO 2 LEVEL 3
46 A PIKAA FVAWITLYGS EX ARO 1 LEVEL 3 EX ARO 2 LEVEL 3
47 A PIKAA NVATSQ EX ARO 1 LEVEL 3 EX ARO 2 LEVEL 3
48 A PIKAA NVAKTSRQ EX ARO 1 LEVEL 3 EX ARO 2 LEVEL 3
49 A PIKAA FVAWITLYGS EX ARO 1 LEVEL 3 EX ARO 2 LEVEL 3
50 A PIKAA NVAETSDQ EX ARO 1 LEVEL 3 EX ARO 2 LEVEL 3
51 A PIKAA NVATSQ EX ARO 1 LEVEL 3 EX ARO 2 LEVEL 3
52 A PIKAA FVAWITLYGS EX ARO 1 LEVEL 3 EX ARO 2 LEVEL 3
53 A PIKAA FVAWITLYGS EX ARO 1 LEVEL 3 EX ARO 2 LEVEL 3
54 A PIKAA NVATSQ EX ARO 1 LEVEL 3 EX ARO 2 LEVEL 3
55 A PIKAA NVAKTSRQ EX ARO 1 LEVEL 3 EX ARO 2 LEVEL 3
56 A PIKAA FVAWITLYGS EX ARO 1 LEVEL 3 EX ARO 2 LEVEL 3
57 A PIKAA NVFAWEIKTLYSDRQ EX ARO 1 LEVEL 3 EX ARO 2 LEVEL 3
58 A PIKAA NVAETSDQ EX ARO 1 LEVEL 3 EX ARO 2 LEVEL 3
59 A PIKAA NVAEKTSDRQ EX ARO 1 LEVEL 3 EX ARO 2 LEVEL 3
60 A PIKAA NVFAWEIKTLYSDRQ EX ARO 1 LEVEL 3 EX ARO 2 LEVEL 3
63 A PIKAA NVAETSDQ EX ARO 1 LEVEL 3 EX ARO 2 LEVEL 3
64 A PIKAA FVAWITLYGS EX ARO 1 LEVEL 3 EX ARO 2 LEVEL 3
65 A PIKAA FVAWITLYGS EX ARO 1 LEVEL 3 EX ARO 2 LEVEL 3
66 A PIKAA NVAKTSRQ EX ARO 1 LEVEL 3 EX ARO 2 LEVEL 3
67 A PIKAA NVAKTSRQ EX ARO 1 LEVEL 3 EX ARO 2 LEVEL 3
68 A PIKAA G EX ARO 1 LEVEL 3 EX ARO 2 LEVEL 3
69 A PIKAA NVAETSDQ EX ARO 1 LEVEL 3 EX ARO 2 LEVEL 3
70 A PIKAA NVAETSDQ EX ARO 1 LEVEL 3 EX ARO 2 LEVEL 3
71 A PIKAA FVAWITLYGSNQ EX ARO 1 LEVEL 3 EX ARO 2 LEVEL 3

72 A PIKAA FVAWITLYGS EX ARO 1 LEVEL 3 EX ARO 2 LEVEL 3
73 A PIKAA NVATSQ EX ARO 1 LEVEL 3 EX ARO 2 LEVEL 3
74 A PIKAA NVAKTSRQ EX ARO 1 LEVEL 3 EX ARO 2 LEVEL 3
76 A PIKAA NVAKTSRQ EX ARO 1 LEVEL 3 EX ARO 2 LEVEL 3
77 A PIKAA NVAETSDQ EX ARO 1 LEVEL 3 EX ARO 2 LEVEL 3
78 A PIKAA FVAWITLYGSNQ EX ARO 1 LEVEL 3 EX ARO 2 LEVEL 3
79 A PIKAA FVAWITLYGS EX ARO 1 LEVEL 3 EX ARO 2 LEVEL 3
80 A PIKAA NVATSQ EX ARO 1 LEVEL 3 EX ARO 2 LEVEL 3
81 A PIKAA NVAETSDQ EX ARO 1 LEVEL 3 EX ARO 2 LEVEL 3
82 A PIKAA FVAWITLYGS EX ARO 1 LEVEL 3 EX ARO 2 LEVEL 3
83 A PIKAA NVAKTSRQ EX ARO 1 LEVEL 3 EX ARO 2 LEVEL 3
84 A PIKAA NVAETSDQ EX ARO 1 LEVEL 3 EX ARO 2 LEVEL 3
85 A PIKAA FVAWITLYGS EX ARO 1 LEVEL 3 EX ARO 2 LEVEL 3
86 A PIKAA FVAWITLYGS EX ARO 1 LEVEL 3 EX ARO 2 LEVEL 3
87 A PIKAA NVAKTSRQ EX ARO 1 LEVEL 3 EX ARO 2 LEVEL 3
88 A PIKAA NVAETGSDQ EX ARO 1 LEVEL 3 EX ARO 2 LEVEL 3
91 A PIKAA NVAEKTSDRQ EX ARO 1 LEVEL 3 EX ARO 2 LEVEL 3
92 A PIKAA NVAETSDQ EX ARO 1 LEVEL 3 EX ARO 2 LEVEL 3
93 A PIKAA NVAKTSRQ EX ARO 1 LEVEL 3 EX ARO 2 LEVEL 3
94 A PIKAA FVAWITLYGS EX ARO 1 LEVEL 3 EX ARO 2 LEVEL 3
95 A PIKAA NVAKTSRQ EX ARO 1 LEVEL 3 EX ARO 2 LEVEL 3
96 A PIKAA NVATSQ EX ARO 1 LEVEL 3 EX ARO 2 LEVEL 3
97 A PIKAA FVAWITLYGS EX ARO 1 LEVEL 3 EX ARO 2 LEVEL 3
98 A PIKAA FVAWITLYGS EX ARO 1 LEVEL 3 EX ARO 2 LEVEL 3
99 A PIKAA NVAETSDQ EX ARO 1 LEVEL 3 EX ARO 2 LEVEL 3
100 A PIKAA NVAETSDQ EX ARO 1 LEVEL 3 EX ARO 2 LEVEL 3
101 A PIKAA FVAWITLYGS EX ARO 1 LEVEL 3 EX ARO 2 LEVEL 3
102 A PIKAA NVAKTSRQ EX ARO 1 LEVEL 3 EX ARO 2 LEVEL 3
103 A PIKAA NVATSQ EX ARO 1 LEVEL 3 EX ARO 2 LEVEL 3
104 A PIKAA N EX ARO 1 LEVEL 3 EX ARO 2 LEVEL 3
105 A PIKAA FVAWITLYGS EX ARO 1 LEVEL 3 EX ARO 2 LEVEL 3
106 A PIKAA NVATSQ EX ARO 1 LEVEL 3 EX ARO 2 LEVEL 3
107 A PIKAA NVAETSDQ EX ARO 1 LEVEL 3 EX ARO 2 LEVEL 3
108 A PIKAA FVAWITLYGS EX ARO 1 LEVEL 3 EX ARO 2 LEVEL 3
109 A PIKAA NVAKTSRQ EX ARO 1 LEVEL 3 EX ARO 2 LEVEL 3
110 A PIKAA NVATSQ EX ARO 1 LEVEL 3 EX ARO 2 LEVEL 3
111 A PIKAA FVAWITLYGS EX ARO 1 LEVEL 3 EX ARO 2 LEVEL 3
112 A PIKAA FVAWITLYGS EX ARO 1 LEVEL 3 EX ARO 2 LEVEL 3
113 A PIKAA NVATSQ EX ARO 1 LEVEL 3 EX ARO 2 LEVEL 3
114 A PIKAA NVAETSDQ EX ARO 1 LEVEL 3 EX ARO 2 LEVEL 3
115 A PIKAA FVAWITLYGS EX ARO 1 LEVEL 3 EX ARO 2 LEVEL 3
116 A PIKAA NVAETSDQ EX ARO 1 LEVEL 3 EX ARO 2 LEVEL 3
117 A PIKAA NVAKTSRQ EX ARO 1 LEVEL 3 EX ARO 2 LEVEL 3
118 A PIKAA NVAEKTSDRQ EX ARO 1 LEVEL 3 EX ARO 2 LEVEL 3
119 A PIKAA NVFAWEIKLYGSDRQ EX ARO 1 LEVEL 3 EX ARO 2 LEVEL 3

**resfile_lvl1.txt**

```
NATRO
ALLAAxc
NOTAA MTC
USE_INPUT_SC
start

16 A PIKAA N EX 1 LEVEL 1 EX 2 LEVEL 1
75 A PIKAA N EX 1 LEVEL 1 EX 2 LEVEL 1

30 A PIKAA G EX 1 LEVEL 1 EX 2 LEVEL 1
31 A PIKAA D EX 1 LEVEL 1 EX 2 LEVEL 1

61 A PIKAA G EX 1 LEVEL 1 EX 2 LEVEL 1
62 A PIKAA D EX 1 LEVEL 1 EX 2 LEVEL 1

89 A PIKAA G EX 1 LEVEL 1 EX 2 LEVEL 1
90 A PIKAA D EX 1 LEVEL 1 EX 2 LEVEL 1

1 A PIKAA S EX 1 LEVEL 1 EX 2 LEVEL 1
2 A PIKAA NVAEKTSDRQ EX 1 LEVEL 1 EX 2 LEVEL 1
3 A PIKAA NVATSQRK EX 1 LEVEL 1 EX 2 LEVEL 1
4 A PIKAA NVATSQ EX 1 LEVEL 1 EX 2 LEVEL 1
5 A PIKAA NVAETSDQ EX 1 LEVEL 1 EX 2 LEVEL 1
6 A PIKAA FVAWITLYGS EX 1 LEVEL 1 EX 2 LEVEL 1
7 A PIKAA NVATSQ EX 1 LEVEL 1 EX 2 LEVEL 1
8 A PIKAA NVAETSDQ EX 1 LEVEL 1 EX 2 LEVEL 1
9 A PIKAA G EX 1 LEVEL 1 EX 2 LEVEL 1
10 A PIKAA NVAKTSRQ EX 1 LEVEL 1 EX 2 LEVEL 1
11 A PIKAA NVAKTSRQ EX 1 LEVEL 1 EX 2 LEVEL 1
12 A PIKAA FVAWITLYGSNQ EX 1 LEVEL 1 EX 2 LEVEL 1
13 A PIKAA W EX 1 LEVEL 1 EX 2 LEVEL 1
14 A PIKAA NVATSQ EX 1 LEVEL 1 EX 2 LEVEL 1
15 A PIKAA NVAETSDQ EX 1 LEVEL 1 EX 2 LEVEL 1
17 A PIKAA NVAETSDQ EX 1 LEVEL 1 EX 2 LEVEL 1
18 A PIKAA NVAKTSRQ EX 1 LEVEL 1 EX 2 LEVEL 1
19 A PIKAA NVFAWEIKTLYSDRQ EX 1 LEVEL 1 EX 2 LEVEL 1
20 A PIKAA W EX 1 LEVEL 1 EX 2 LEVEL 1
21 A PIKAA NVATSQ EX 1 LEVEL 1 EX 2 LEVEL 1
22 A PIKAA NVAKTSRQ EX 1 LEVEL 1 EX 2 LEVEL 1
23 A PIKAA FVAWITLYGS EX 1 LEVEL 1 EX 2 LEVEL 1
24 A PIKAA NVAETSDQ EX 1 LEVEL 1 EX 2 LEVEL 1
25 A PIKAA NVATSQ EX 1 LEVEL 1 EX 2 LEVEL 1
26 A PIKAA NVAKTSRQ EX 1 LEVEL 1 EX 2 LEVEL 1
27 A PIKAA FVAWITLYGS EX 1 LEVEL 1 EX 2 LEVEL 1
```

28 A PIKAA NVAETSDQ EX 1 LEVEL 1 EX 2 LEVEL 1
29 A PIKAA ST EX 1 LEVEL 1 EX 2 LEVEL 1
31 A PIKAA NVAEKTGSDRQ EX 1 LEVEL 1 EX 2 LEVEL 1
32 A PIKAA NVAEKTSDRQ EX 1 LEVEL 1 EX 2 LEVEL 1
33 A PIKAA NVATSQ EX 1 LEVEL 1 EX 2 LEVEL 1
34 A PIKAA NVAETSDQ EX 1 LEVEL 1 EX 2 LEVEL 1
35 A PIKAA FVAWITLYGS EX 1 LEVEL 1 EX 2 LEVEL 1
36 A PIKAA NVAETSDQ EX 1 LEVEL 1 EX 2 LEVEL 1
37 A PIKAA NVATSQ EX 1 LEVEL 1 EX 2 LEVEL 1
38 A PIKAA FVAWITLYGS EX 1 LEVEL 1 EX 2 LEVEL 1
39 A PIKAA FVAWITLYGS EX 1 LEVEL 1 EX 2 LEVEL 1
40 A PIKAA NVAKTSRQ EX 1 LEVEL 1 EX 2 LEVEL 1
41 A PIKAA NVAKTSRQ EX 1 LEVEL 1 EX 2 LEVEL 1
42 A PIKAA FVAWITLYGS EX 1 LEVEL 1 EX 2 LEVEL 1
43 A PIKAA NVAETSDQ EX 1 LEVEL 1 EX 2 LEVEL 1
44 A PIKAA NVATSQ EX 1 LEVEL 1 EX 2 LEVEL 1
45 A PIKAA N EX 1 LEVEL 1 EX 2 LEVEL 1
46 A PIKAA FVAWITLYGS EX 1 LEVEL 1 EX 2 LEVEL 1
47 A PIKAA NVATSQ EX 1 LEVEL 1 EX 2 LEVEL 1
48 A PIKAA NVAKTSRQ EX 1 LEVEL 1 EX 2 LEVEL 1
49 A PIKAA FVAWITLYGS EX 1 LEVEL 1 EX 2 LEVEL 1
50 A PIKAA NVAETSDQ EX 1 LEVEL 1 EX 2 LEVEL 1
51 A PIKAA NVATSQ EX 1 LEVEL 1 EX 2 LEVEL 1
52 A PIKAA FVAWITLYGS EX 1 LEVEL 1 EX 2 LEVEL 1
53 A PIKAA FVAWITLYGS EX 1 LEVEL 1 EX 2 LEVEL 1
54 A PIKAA NVATSQ EX 1 LEVEL 1 EX 2 LEVEL 1
55 A PIKAA NVAKTSRQ EX 1 LEVEL 1 EX 2 LEVEL 1
56 A PIKAA FVAWITLYGS EX 1 LEVEL 1 EX 2 LEVEL 1
57 A PIKAA NVFAWEIKTLYSDRQ EX 1 LEVEL 1 EX 2 LEVEL 1
58 A PIKAA NVAETSDQ EX 1 LEVEL 1 EX 2 LEVEL 1
59 A PIKAA NVAEKTSDRQ EX 1 LEVEL 1 EX 2 LEVEL 1
60 A PIKAA NVFAWEIKTLYSDRQ EX 1 LEVEL 1 EX 2 LEVEL 1
63 A PIKAA NVAETSDQ EX 1 LEVEL 1 EX 2 LEVEL 1
64 A PIKAA FVAWITLYGS EX 1 LEVEL 1 EX 2 LEVEL 1
65 A PIKAA FVAWITLYGS EX 1 LEVEL 1 EX 2 LEVEL 1
66 A PIKAA NVAKTSRQ EX 1 LEVEL 1 EX 2 LEVEL 1
67 A PIKAA NVAKTSRQ EX 1 LEVEL 1 EX 2 LEVEL 1
68 A PIKAA G EX 1 LEVEL 1 EX 2 LEVEL 1
69 A PIKAA NVAETSDQ EX 1 LEVEL 1 EX 2 LEVEL 1
70 A PIKAA NVAETSDQ EX 1 LEVEL 1 EX 2 LEVEL 1
71 A PIKAA FVAWITLYGSNQ EX 1 LEVEL 1 EX 2 LEVEL 1
72 A PIKAA FVAWITLYGS EX 1 LEVEL 1 EX 2 LEVEL 1
73 A PIKAA NVATSQ EX 1 LEVEL 1 EX 2 LEVEL 1
74 A PIKAA NVAKTSRQ EX 1 LEVEL 1 EX 2 LEVEL 1
76 A PIKAA NVAKTSRQ EX 1 LEVEL 1 EX 2 LEVEL 1
77 A PIKAA NVAETSDQ EX 1 LEVEL 1 EX 2 LEVEL 1

78 A PIKAA FVAWITLYGSNQ EX 1 LEVEL 1 EX 2 LEVEL 1
79 A PIKAA FVAWITLYGS EX 1 LEVEL 1 EX 2 LEVEL 1
80 A PIKAA NVATSQ EX 1 LEVEL 1 EX 2 LEVEL 1
81 A PIKAA NVAETSDQ EX 1 LEVEL 1 EX 2 LEVEL 1
82 A PIKAA FVAWITLYGS EX 1 LEVEL 1 EX 2 LEVEL 1
83 A PIKAA NVAKTSRQ EX 1 LEVEL 1 EX 2 LEVEL 1
84 A PIKAA NVAETSDQ EX 1 LEVEL 1 EX 2 LEVEL 1
85 A PIKAA FVAWITLYGS EX 1 LEVEL 1 EX 2 LEVEL 1
86 A PIKAA FVAWITLYGS EX 1 LEVEL 1 EX 2 LEVEL 1
87 A PIKAA NVAKTSRQ EX 1 LEVEL 1 EX 2 LEVEL 1
88 A PIKAA NVAETGSDQ EX 1 LEVEL 1 EX 2 LEVEL 1
91 A PIKAA NVAEKTSDRQ EX 1 LEVEL 1 EX 2 LEVEL 1
92 A PIKAA NVAETSDQ EX 1 LEVEL 1 EX 2 LEVEL 1
93 A PIKAA NVAKTSRQ EX 1 LEVEL 1 EX 2 LEVEL 1
94 A PIKAA FVAWITLYGS EX 1 LEVEL 1 EX 2 LEVEL 1
95 A PIKAA NVAKTSRQ EX 1 LEVEL 1 EX 2 LEVEL 1
96 A PIKAA NVATSQ EX 1 LEVEL 1 EX 2 LEVEL 1
97 A PIKAA FVAWITLYGS EX 1 LEVEL 1 EX 2 LEVEL 1
98 A PIKAA FVAWITLYGS EX 1 LEVEL 1 EX 2 LEVEL 1
99 A PIKAA NVAETSDQ EX 1 LEVEL 1 EX 2 LEVEL 1
100 A PIKAA NVAETSDQ EX 1 LEVEL 1 EX 2 LEVEL 1
101 A PIKAA FVAWITLYGS EX 1 LEVEL 1 EX 2 LEVEL 1
102 A PIKAA NVAKTSRQ EX 1 LEVEL 1 EX 2 LEVEL 1
103 A PIKAA NVATSQ EX 1 LEVEL 1 EX 2 LEVEL 1
104 A PIKAA N EX 1 LEVEL 1 EX 2 LEVEL 1
105 A PIKAA FVAWITLYGS EX 1 LEVEL 1 EX 2 LEVEL 1
106 A PIKAA NVATSQ EX 1 LEVEL 1 EX 2 LEVEL 1
107 A PIKAA NVAETSDQ EX 1 LEVEL 1 EX 2 LEVEL 1
108 A PIKAA FVAWITLYGS EX 1 LEVEL 1 EX 2 LEVEL 1
109 A PIKAA NVAKTSRQ EX 1 LEVEL 1 EX 2 LEVEL 1
110 A PIKAA NVATSQ EX 1 LEVEL 1 EX 2 LEVEL 1
111 A PIKAA FVAWITLYGS EX 1 LEVEL 1 EX 2 LEVEL 1
112 A PIKAA FVAWITLYGS EX 1 LEVEL 1 EX 2 LEVEL 1
113 A PIKAA NVATSQ EX 1 LEVEL 1 EX 2 LEVEL 1
114 A PIKAA NVAETSDQ EX 1 LEVEL 1 EX 2 LEVEL 1
115 A PIKAA FVAWITLYGS EX 1 LEVEL 1 EX 2 LEVEL 1
116 A PIKAA NVAETSDQ EX 1 LEVEL 1 EX 2 LEVEL 1
117 A PIKAA NVAKTSRQ EX 1 LEVEL 1 EX 2 LEVEL 1
118 A PIKAA NVAEKTSDRQ EX 1 LEVEL 1 EX 2 LEVEL 1
119 A PIKAA NVFAWEIKLYGSDRQ EX 1 LEVEL 1 EX 2 LEVEL 1

**resfile_lvl3.txt**

NATRO
ALLAAxc
NOTAA MTC

USE_INPUT_SC
start

16 A PIKAA N EX 1 LEVEL 3 EX 2 LEVEL 3
75 A PIKAA N EX 1 LEVEL 3 EX 2 LEVEL 3

30 A PIKAA G EX 1 LEVEL 3 EX 2 LEVEL 3
31 A PIKAA D EX 1 LEVEL 3 EX 2 LEVEL 3

61 A PIKAA G EX 1 LEVEL 3 EX 2 LEVEL 3
62 A PIKAA D EX 1 LEVEL 3 EX 2 LEVEL 3

89 A PIKAA G EX 1 LEVEL 3 EX 2 LEVEL 3
90 A PIKAA D EX 1 LEVEL 3 EX 2 LEVEL 3

1 A PIKAA S EX 1 LEVEL 3 EX 2 LEVEL 3
2 A PIKAA NVAEKTSDRQ EX 1 LEVEL 3 EX 2 LEVEL 3
3 A PIKAA NVATSQRK EX 1 LEVEL 3 EX 2 LEVEL 3
4 A PIKAA NVATSQ EX 1 LEVEL 3 EX 2 LEVEL 3
5 A PIKAA NVAETSDQ EX 1 LEVEL 3 EX 2 LEVEL 3
6 A PIKAA FVAWITLYGS EX 1 LEVEL 3 EX 2 LEVEL 3
7 A PIKAA NVATSQ EX 1 LEVEL 3 EX 2 LEVEL 3
8 A PIKAA NVAETSDQ EX 1 LEVEL 3 EX 2 LEVEL 3
9 A PIKAA G EX 1 LEVEL 3 EX 2 LEVEL 3
10 A PIKAA NVAKTSRQ EX 1 LEVEL 3 EX 2 LEVEL 3
11 A PIKAA NVAKTSRQ EX 1 LEVEL 3 EX 2 LEVEL 3
12 A PIKAA FVAWITLYGSNQ EX 1 LEVEL 3 EX 2 LEVEL 3
13 A PIKAA W EX 1 LEVEL 3 EX 2 LEVEL 3
14 A PIKAA NVATSQ EX 1 LEVEL 3 EX 2 LEVEL 3
15 A PIKAA NVAETSDQ EX 1 LEVEL 3 EX 2 LEVEL 3
17 A PIKAA NVAETSDQ EX 1 LEVEL 3 EX 2 LEVEL 3
18 A PIKAA NVAKTSRQ EX 1 LEVEL 3 EX 2 LEVEL 3
19 A PIKAA NVFAWEIKTLYSDRQ EX 1 LEVEL 3 EX 2 LEVEL 3
20 A PIKAA W EX 1 LEVEL 3 EX 2 LEVEL 3
21 A PIKAA NVATSQ EX 1 LEVEL 3 EX 2 LEVEL 3
22 A PIKAA NVAKTSRQ EX 1 LEVEL 3 EX 2 LEVEL 3
23 A PIKAA FVAWITLYGS EX 1 LEVEL 3 EX 2 LEVEL 3
24 A PIKAA NVAETSDQ EX 1 LEVEL 3 EX 2 LEVEL 3
25 A PIKAA NVATSQ EX 1 LEVEL 3 EX 2 LEVEL 3
26 A PIKAA NVAKTSRQ EX 1 LEVEL 3 EX 2 LEVEL 3
27 A PIKAA FVAWITLYGS EX 1 LEVEL 3 EX 2 LEVEL 3
28 A PIKAA NVAETSDQ EX 1 LEVEL 3 EX 2 LEVEL 3
29 A PIKAA ST EX 1 LEVEL 3 EX 2 LEVEL 3
31 A PIKAA NVAEKTGSDRQ EX 1 LEVEL 3 EX 2 LEVEL 3
32 A PIKAA NVAEKTSDRQ EX 1 LEVEL 3 EX 2 LEVEL 3
33 A PIKAA NVATSQ EX 1 LEVEL 3 EX 2 LEVEL 3

34 A PIKAA NVAETSDQ EX 1 LEVEL 3 EX 2 LEVEL 3
35 A PIKAA FVAWITLYGS EX 1 LEVEL 3 EX 2 LEVEL 3
36 A PIKAA NVAETSDQ EX 1 LEVEL 3 EX 2 LEVEL 3
37 A PIKAA NVATSQ EX 1 LEVEL 3 EX 2 LEVEL 3
38 A PIKAA FVAWITLYGS EX 1 LEVEL 3 EX 2 LEVEL 3
39 A PIKAA FVAWITLYGS EX 1 LEVEL 3 EX 2 LEVEL 3
40 A PIKAA NVAKTSRQ EX 1 LEVEL 3 EX 2 LEVEL 3
41 A PIKAA NVAKTSRQ EX 1 LEVEL 3 EX 2 LEVEL 3
42 A PIKAA FVAWITLYGS EX 1 LEVEL 3 EX 2 LEVEL 3
43 A PIKAA NVAETSDQ EX 1 LEVEL 3 EX 2 LEVEL 3
44 A PIKAA NVATSQ EX 1 LEVEL 3 EX 2 LEVEL 3
45 A PIKAA N EX 1 LEVEL 3 EX 2 LEVEL 3
46 A PIKAA FVAWITLYGS EX 1 LEVEL 3 EX 2 LEVEL 3
47 A PIKAA NVATSQ EX 1 LEVEL 3 EX 2 LEVEL 3
48 A PIKAA NVAKTSRQ EX 1 LEVEL 3 EX 2 LEVEL 3
49 A PIKAA FVAWITLYGS EX 1 LEVEL 3 EX 2 LEVEL 3
50 A PIKAA NVAETSDQ EX 1 LEVEL 3 EX 2 LEVEL 3
51 A PIKAA NVATSQ EX 1 LEVEL 3 EX 2 LEVEL 3
52 A PIKAA FVAWITLYGS EX 1 LEVEL 3 EX 2 LEVEL 3
53 A PIKAA FVAWITLYGS EX 1 LEVEL 3 EX 2 LEVEL 3
54 A PIKAA NVATSQ EX 1 LEVEL 3 EX 2 LEVEL 3
55 A PIKAA NVAKTSRQ EX 1 LEVEL 3 EX 2 LEVEL 3
56 A PIKAA FVAWITLYGS EX 1 LEVEL 3 EX 2 LEVEL 3
57 A PIKAA NVFAWEIKTLYSDRQ EX 1 LEVEL 3 EX 2 LEVEL 3
58 A PIKAA NVAETSDQ EX 1 LEVEL 3 EX 2 LEVEL 3
59 A PIKAA NVAEKTSDRQ EX 1 LEVEL 3 EX 2 LEVEL 3
60 A PIKAA NVFAWEIKTLYSDRQ EX 1 LEVEL 3 EX 2 LEVEL 3
63 A PIKAA NVAETSDQ EX 1 LEVEL 3 EX 2 LEVEL 3
64 A PIKAA FVAWITLYGS EX 1 LEVEL 3 EX 2 LEVEL 3
65 A PIKAA FVAWITLYGS EX 1 LEVEL 3 EX 2 LEVEL 3
66 A PIKAA NVAKTSRQ EX 1 LEVEL 3 EX 2 LEVEL 3
67 A PIKAA NVAKTSRQ EX 1 LEVEL 3 EX 2 LEVEL 3
68 A PIKAA G EX 1 LEVEL 3 EX 2 LEVEL 3
69 A PIKAA NVAETSDQ EX 1 LEVEL 3 EX 2 LEVEL 3
70 A PIKAA NVAETSDQ EX 1 LEVEL 3 EX 2 LEVEL 3
71 A PIKAA FVAWITLYGSNQ EX 1 LEVEL 3 EX 2 LEVEL 3
72 A PIKAA FVAWITLYGS EX 1 LEVEL 3 EX 2 LEVEL 3
73 A PIKAA NVATSQ EX 1 LEVEL 3 EX 2 LEVEL 3
74 A PIKAA NVAKTSRQ EX 1 LEVEL 3 EX 2 LEVEL 3
76 A PIKAA NVAKTSRQ EX 1 LEVEL 3 EX 2 LEVEL 3
77 A PIKAA NVAETSDQ EX 1 LEVEL 3 EX 2 LEVEL 3
78 A PIKAA FVAWITLYGSNQ EX 1 LEVEL 3 EX 2 LEVEL 3
79 A PIKAA FVAWITLYGS EX 1 LEVEL 3 EX 2 LEVEL 3
80 A PIKAA NVATSQ EX 1 LEVEL 3 EX 2 LEVEL 3
81 A PIKAA NVAETSDQ EX 1 LEVEL 3 EX 2 LEVEL 3
82 A PIKAA FVAWITLYGS EX 1 LEVEL 3 EX 2 LEVEL 3

```
83 A PIKAA NVAKTSRQ EX 1 LEVEL 3 EX 2 LEVEL 3
84 A PIKAA NVAETSDQ EX 1 LEVEL 3 EX 2 LEVEL 3
85 A PIKAA FVAWITLYGS EX 1 LEVEL 3 EX 2 LEVEL 3
86 A PIKAA FVAWITLYGS EX 1 LEVEL 3 EX 2 LEVEL 3
87 A PIKAA NVAKTSRQ EX 1 LEVEL 3 EX 2 LEVEL 3
88 A PIKAA NVAETGSDQ EX 1 LEVEL 3 EX 2 LEVEL 3
91 A PIKAA NVAEKTSDRQ EX 1 LEVEL 3 EX 2 LEVEL 3
92 A PIKAA NVAETSDQ EX 1 LEVEL 3 EX 2 LEVEL 3
93 A PIKAA NVAKTSRQ EX 1 LEVEL 3 EX 2 LEVEL 3
94 A PIKAA FVAWITLYGS EX 1 LEVEL 3 EX 2 LEVEL 3
95 A PIKAA NVAKTSRQ EX 1 LEVEL 3 EX 2 LEVEL 3
96 A PIKAA NVATSQ EX 1 LEVEL 3 EX 2 LEVEL 3
97 A PIKAA FVAWITLYGS EX 1 LEVEL 3 EX 2 LEVEL 3
98 A PIKAA FVAWITLYGS EX 1 LEVEL 3 EX 2 LEVEL 3
99 A PIKAA NVAETSDQ EX 1 LEVEL 3 EX 2 LEVEL 3
100 A PIKAA NVAETSDQ EX 1 LEVEL 3 EX 2 LEVEL 3
101 A PIKAA FVAWITLYGS EX 1 LEVEL 3 EX 2 LEVEL 3
102 A PIKAA NVAKTSRQ EX 1 LEVEL 3 EX 2 LEVEL 3
103 A PIKAA NVATSQ EX 1 LEVEL 3 EX 2 LEVEL 3
104 A PIKAA N EX 1 LEVEL 3 EX 2 LEVEL 3
105 A PIKAA FVAWITLYGS EX 1 LEVEL 3 EX 2 LEVEL 3
106 A PIKAA NVATSQ EX 1 LEVEL 3 EX 2 LEVEL 3
107 A PIKAA NVAETSDQ EX 1 LEVEL 3 EX 2 LEVEL 3
108 A PIKAA FVAWITLYGS EX 1 LEVEL 3 EX 2 LEVEL 3
109 A PIKAA NVAKTSRQ EX 1 LEVEL 3 EX 2 LEVEL 3
110 A PIKAA NVATSQ EX 1 LEVEL 3 EX 2 LEVEL 3
111 A PIKAA FVAWITLYGS EX 1 LEVEL 3 EX 2 LEVEL 3
112 A PIKAA FVAWITLYGS EX 1 LEVEL 3 EX 2 LEVEL 3
113 A PIKAA NVATSQ EX 1 LEVEL 3 EX 2 LEVEL 3
114 A PIKAA NVAETSDQ EX 1 LEVEL 3 EX 2 LEVEL 3
115 A PIKAA FVAWITLYGS EX 1 LEVEL 3 EX 2 LEVEL 3
116 A PIKAA NVAETSDQ EX 1 LEVEL 3 EX 2 LEVEL 3
117 A PIKAA NVAKTSRQ EX 1 LEVEL 3 EX 2 LEVEL 3
118 A PIKAA NVAEKTSDRQ EX 1 LEVEL 3 EX 2 LEVEL 3
119 A PIKAA NVFAWEIKLYGSDRQ EX 1 LEVEL 3 EX 2 LEVEL 3
```

**rscript_flexbb_relax.xml**

```
<ROSETTASCRIPTS>
	<SCOREFXNS>
		<ScoreFunction name="cstscore" weights="ref2015">
			<Reweight scoretype="aa_composition" weight="1.0"/>
			<Reweight scoretype="netcharge" weight="1.0" />
			<Reweight scoretype="atom_pair_constraint" weight="1"/>
			<Reweight scoretype="angle_constraint" weight="1"/>
			<Reweight scoretype="dihedral_constraint" weight="1"/>
```

```
				<Set aa_composition_setup_file="trp_ala_thr_ser_asn.comp" />
				<Set netcharge_setup_file="netcharge.charge" />
			</ScoreFunction>
			<ScoreFunction name="cstscore_soft" weights="ref2015">
				<Reweight scoretype="aa_composition" weight="1.0"/>
				<Reweight scoretype="netcharge" weight="1.0" />
				<Reweight scoretype="atom_pair_constraint" weight="1"/>
				<Reweight scoretype="angle_constraint" weight="1"/>
				<Reweight scoretype="dihedral_constraint" weight="1"/>
				<Reweight scoretype="fa_rep" weight="0.2"/>
				<Set aa_composition_setup_file="trp_ala_thr_ser_asn.comp" />
				<Set netcharge_setup_file="netcharge.charge" />
			</ScoreFunction>
			<ScoreFunction name="score" weights="ref2015"/>
		</SCOREFXNS>
		<TASKOPERATIONS>
			<ReadResfile name="rr_ex" filename="resfile_topology_lvl1_round28.txt"/>
			<ReadResfile name="rr_aro" filename="resfile_topology_aro_round28.txt"/>
			<ReadResfile name="rr_ex_3" filename="resfile_topology_lvl3_round28.txt"/>
			<InitializeFromCommandline name="ifcl" />
		</TASKOPERATIONS>
		<FILTERS>
			<PackStat name="pstat_mc" threshold="0.48" repeats="10"/>
			<PackStat name="pstat_stop" threshold="0.65" repeats="10"/>
			<PackStat name="pstat" repeats="10" confidence="0"/>
			<NetCharge name="net_charge" confidence="0"/>
		</FILTERS>
		<MOVERS>
			<ConstraintSetMover name="csts" add_constraints="1"
cst_file="constraints_NDI_122021.cst"/>
			<PackRotamersMover name="fixbb_aro" scorefxn="cstscore_soft"
task_operations="rr_aro,ifcl"/>
			<PackRotamersMover name="fixbb_ex_1" scorefxn="cstscore"
task_operations="rr_ex,ifcl"/>
			<PackRotamersMover name="fixbb_ex_3" scorefxn="cstscore"
task_operations="rr_ex_3,ifcl"/>
			<MinMover name="min_bb" scorefxn="cstscore" tolerance="0.000001"
max_iter="10000" chi="0" bb="1"/>
			<MinMover name="min_sc" scorefxn="cstscore" tolerance="0.000001"
max_iter="10000" chi="1" bb="0"/>
			<FastRelax name="fr" scorefxn="cstscore" relaxscript="KillA2019" />
			<ParsedProtocol name="custom_flex">
				<Add mover_name="fixbb_ex_1"/>
				<Add mover_name="fr"/>
			</ParsedProtocol>
			<ParsedProtocol name="final_flex">
```

```
				<Add mover_name="fixbb_ex_3"/>
				<Add mover_name="fr"/>
			</ParsedProtocol>
			<GenericMonteCarlo name="iterate" mover_name="custom_flex"
							filter_name="pstat_mc" trials="3" preapply="0"
							temperature="0" sample_type="high"
stopping_condition="pstat_stop"
							saved_accept_file_name="last_accepted.pdb"
recover_low="1"/>
			<GenericMonteCarlo name="final" mover_name="final_flex"
							filter_name="pstat_mc" trials="1" preapply="0"
							temperature ="0" sample_type="high"
							saved_accept_file_name="last_accepted.pdb"
recover_low="1"/>
		</MOVERS>
		<APPLY_TO_POSE>
		</APPLY_TO_POSE>
		<PROTOCOLS>
			<Add mover_name="csts"/>
			<Add mover_name="fixbb_aro"/>
			<Add mover_name="min_sc"/>
			<Add mover_name="min_bb"/>
			<Add mover_name="iterate"/>
			<Add mover_name="final"/>
			<Add filter_name="pstat"/>
			<Add filter_name="net_charge"/>
		</PROTOCOLS>
		<OUTPUT scorefxn="cstscore"/>
</ROSETTASCRIPTS>
```

**constraints.cst**

```
#Asn carbonyl Hbond #1
AtomPair ND2 16A O3 1X HARMONIC 3.1 0.2
AtomPair OD1 16A O3 1X HARMONIC 5.2 0.2

#Asn carbonyl Hbond #2
AtomPair ND2 75A O1 1X HARMONIC 2.8 0.2
AtomPair OD1 75A O1 1X HARMONIC 5.0 0.2
```
